# A topography-based predictive framework for naturalistic viewing fMRI

**DOI:** 10.1101/2022.05.26.493420

**Authors:** Xuan Li, Patrick Friedrich, Kaustubh R. Patil, Simon B. Eickhoff, Susanne Weis

## Abstract

Recent work has shown great interest in understanding individual differences in complex brain function under naturalistic viewing (NV) conditions. However, methods specifically designed for achieving this goal remain limited. Here, we propose a novel approach, called TOpography-based Predictive Framework (TOPF), to investigate individual differences in evoked brain activity on NV fMRI data. Specifically, TOPF identifies individual-specific evoked activity topographies in a data-driven manner and examines their behavioural relevance using a machine learning predictive framework. Our results show that these topographies successfully predict individual phenotypes across cognition, emotion and personality on unseen subjects, and the identified predictive brain regions are neurobiologically interpretable. Further, the prediction accuracy exceeds that of the commonly-used functional connectivity-based features. Conceptually, we highlight the importance of examining multivariate evoked activity patterns for studying brain-behaviour relationships. In summary, we provide a powerful tool for understanding individual differences and brain-behaviour relationships on NV fMRI data.

## Introduction

Understanding individual differences in brain function has attracted increasing attention in recent decades [*1*]. Meanwhile, naturalistic viewing (NV) settings, e.g., watching movies in the scanner, have emerged as a promising tool for studying brain function. NV settings use naturalistic stimuli (e.g., movie clips) to approximate real-life situations, overcoming the limited ecological validity of conventional task-based paradigms [*2*]. On the other hand, they improve subject compliance and engagement, overcoming the unsystematic noise caused by unconstrained brain states in resting-state (RS) settings [*3*]. Furthermore, naturalistic stimuli elicit complex cognitive processes that may not be observable when using simplified conventional tasks, such as hierarchical memory in processing unfolding stories [*4*]. While being time-locked across individuals [*5*], brain activity during NV also exhibits substantial individual differences [*6*]. These advantages make NV settings particularly useful for exploring individual differences in higher-order brain functions under naturalistic conditions and ultimately for deepening our understanding of brain-behaviour relationships and psychiatric diseases [*7*]. Yet, methods for a meaningful characterisation of individual differences in brain activity evoked by these complex naturalistic stimuli (i.e., stimulus-evoked activity) remain limited.

Classic approaches such as the general linear model (GLM) have been applied to characterise stimulus-evoked activity in individuals on NV functional magnetic resonance imaging (fMRI) data [*8*]. Such approaches require identification of all features relevant to stimulus-evoked activity (e.g., movie annotations), which is known to be difficult due to the dynamic and multimodal nature of naturalistic stimuli. By contrast, data-driven approaches can detect stimulus-evoked activity without relying on explicit descriptions of the stimulus [*9, 10*]. They leverage the time-locked nature of a stimulus across subjects, typically using the variance shared across subjects to estimate the stimulus-evoked activity and characterise individual differences therein for each region of interest (ROI) or voxel separately [*6*]. Many studies have found associations between these individual differences and individual traits, behaviour and disease status in individual ROIs or voxels [*11, 12*]. While these studies provide valuable insights, our understanding of individual differences could be fundamentally improved by examining spatially distributed brain activity patterns rather than each ROI or voxel separately.

In fact, examining distributed activity patterns facilitates not only the understanding of brain functional organisation but also mapping between brain function and behaviour [*13*–*15*]. Such multivariate patterns will also open the door to a wealth of advanced machine learning models. This will enable behaviour prediction and clinical diagnosis at an individual level and evaluations of the generalisability of inferred brain-behaviour relationships. In turn, it can also provide important insights into the neural substrates of behaviours and biomarkers of diseases [*16, 17*]. While classic approaches for RS fMRI data analysis, such as independent component analysis (ICA) [*18*], can be applied to characterise spatially distributed activity patterns in individuals during NV fMRI, they typically capture the interactions among ROIs or voxels and fail to separate the stimulus-evoked activity from the other fMRI signal components (e.g., spontaneous brain activity). On the other hand, several recently proposed approaches, such as the shared response model (SRM) [*19*] and hyperalignment [*20*], are capable of depicting individual stimulus-evoked activity patterns. Nevertheless, these approaches often focus on the fine-grained patterns across voxels within certain brain regions, with their utility limited by their high memory and computational demands. Therefore, novel approaches that provide a meaningful and efficient characterisation of individual brain activity patterns are required for a better understanding of individual differences on NV fMRI data.

In this study, we propose the TOpography-based Predictive Framework (TOPF) to facilitate understanding of individual differences in brain activity in relation to behaviour on NV fMRI data. TOPF consists of two components: (i) identification of stimulus-evoked activity patterns across all ROIs in individuals based on a whole-brain parcellation in a data-driven way, and (ii) examination of their relationship with individual behaviour via machine learning-based prediction. Specifically, for each ROI separately, TOPF applies a principal component analysis (PCA) to identify stimulus-evoked activity time courses shared across subjects by the detected principal components (PCs). Subject-wise PC loadings thereby reflect the expression levels of these shared time courses specific to each subject. For each subject, a stimulus-evoked activity pattern is characterised by the pattern of these PC loadings across all ROIs, termed individual-specific topography (Fig. 1). These topographies intuitively delineate the unique patterns of how strongly each subject’s brain activity follows the shared stimulus-evoked activity across the whole brain.

**Fig. 1:**
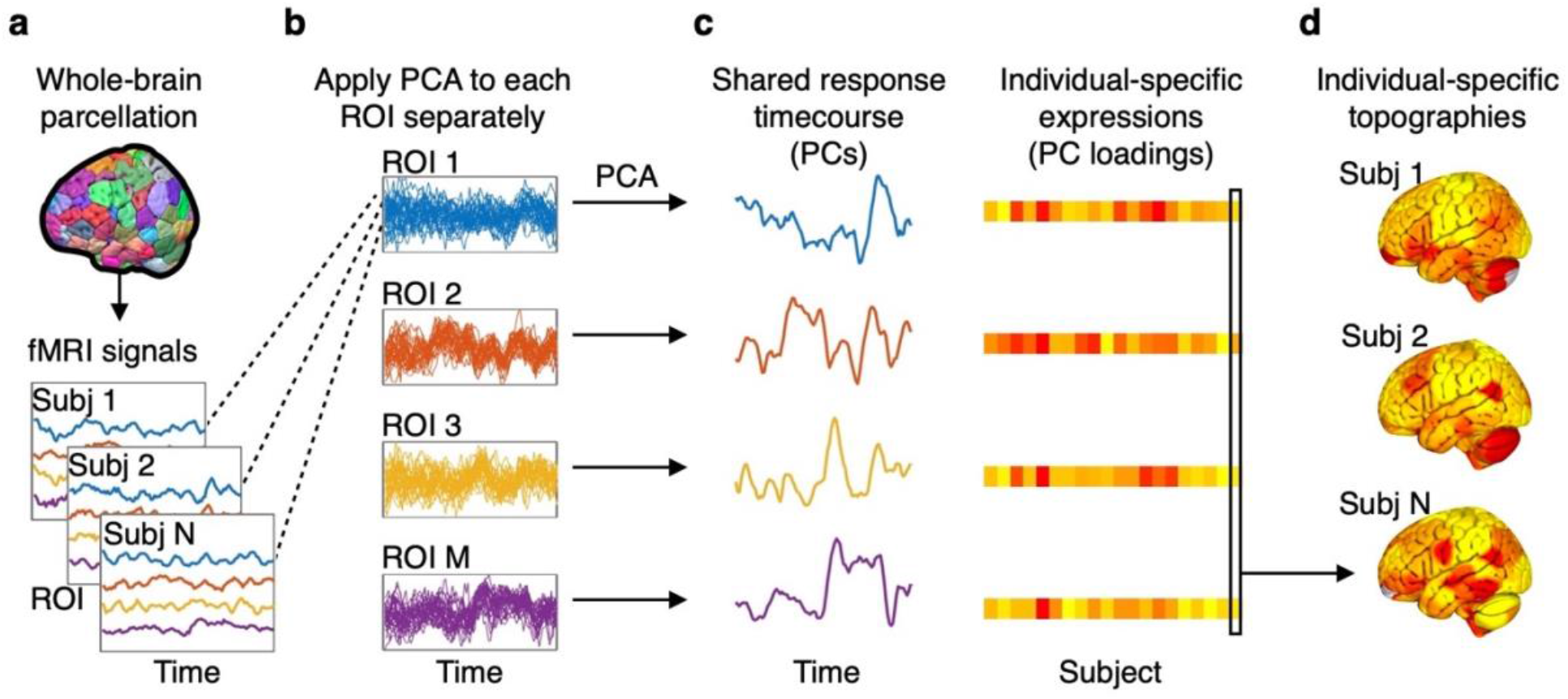
Schematic of TOPF for identifying individual-specific topographies. **a**, For each subject, the whole brain is parcellated into distinct functionally defined ROIs and the fMRI BOLD time series averaged across voxels within each ROI is extracted. **b**, For each ROI, the extracted fMRI time series is z-score standardised, collected across all subjects and then subjected to a PCA. **c**, The resultant PCs and PC loadings of each ROI represent the shared response time series and individual-specific expressions (IEs), respectively. **d**, The pattern of the PC loadings of each subject across all ROIs is defined as an individual-specific topography (e.g., marked by the black box).

Capitalising on fMRI data of multiple NV and task-based paradigms from the human connectome project (HCP) [*21*], we (i) validated the ability and stability of TOPF to capture stimulus-evoked activity, (ii) validated the ability of TOPF to capture meaningful interindividual variability in brain function, (iii) predicted phenotypes across cognition, emotion and personality from individual activity topographies on unseen subjects using a machine learning predictive model, and (iv) demonstrated the promising interpretability of learned predictive models by pinpointing phenotype-related brain regions.

Our approach has several features. First, by highlighting the importance of characterising individual activity topographies, it provides a new perspective to explore individual differences in brain function during NV. Second, by using a cross-validated machine learning framework, it alleviates overfitting when inferring models of brain-behaviour relationships while increasing their generalisability to novel data. Third, it provides a flexible framework that can be applied to not only NV but also conventional task-based fMRI data. Last, it facilitates understanding brain-behaviour relationships by allowing us to localise brain regions that are predictive of behaviours. In sum, TOPF provides a powerful data-driven tool for understanding individual differences and brain-behaviour relationships on NV fMRI data.

## Results

### Identifying individual stimulus-evoked activity patterns

The first step in TOPF is to identify individual stimulus-evoked activity patterns across ROIs. Broadly, the TOPF approach is built on an assumption that each observed NV fMRI time series contains a stimulus-evoked component, such that it is shared but expressed with different intensities across individuals [*6, 22*]. In this study, this time series component and its expression level for each subject are termed shared response and individual-specific expression (IE), respectively. The pattern of the IE values across ROIs for each subject is what we refer to as individual-specific topography. Such individual-specific topographies will be later used as features for phenotype prediction.

Aiming at individual topographies of a reasonable resolution rather than the fine-grained patterns across voxels, we parcellated the whole brain into 268 functionally defined ROIs [*23*] and computed the fMRI time series averaged over voxels within each ROI (Fig. 1a). This reduced the spatial dimensionality of fMRI data, thereby the computational load, and increased the signal-to-noise ratio. TOPF uses PCA to identify the shared response and IEs. For each ROI separately, the voxel-averaged fMRI time series of all subjects were subjected to a PCA after z-score normalisation, where dimensionality reduction was performed on the subject dimension rather than the temporal dimension of the fMRI data (Fig. 1b). The shared response and IEs for each ROI are represented by the detected PC time series and the subject-wise PC loadings, respectively (Fig. 1c). The individual-specific topography of each subject is operationalised as the pattern of that subject’s PC loadings across all ROIs (Fig. 1d). As the first PC (PC1) captures the largest temporal variance shared between subjects across the duration of the scan, thus most likely reflecting the shared response, we will mainly focus on the results of PC1 (see results of PC2 and PC3 in the Supplementary Information).

### Evaluating the validity of TOPF on task-based fMRI data

The validity of TOPF in identifying stimulus-evoked activity was evaluated on a task-based fMRI dataset (dataset1; Supplementary Table 1), a separate dataset from the HCP containing 100 unrelated subjects. Two representative tasks of the seven HCP tasks [*24*] that tapped into two different levels of the hierarchy of brain function, i.e., the motor task and the social cognition (social) task, were used for the evaluation. The availability of the temporal structure of these conventional tasks permitted the application of GLM, which was expected to accurately identify task-evoked activations, thus providing a “ground truth” [*25*].

To evaluate the validity of TOPF-derived activity topographies, for each task, a TOPF-derived group-level topography was compared against a group-level activation (z-score) map derived by GLM [*21*]. The former was defined as the map across ROIs of the variance explained by the shared response (i.e., PC1). The latter was generated by aggregating the activation maps (i.e., computing the maximum absolute values of the z-scores) across all experimental conditions. A high Pearson’s correlation coefficient (*r*) between these two was achieved for both tasks (motor: *r* = 0.72; Fig. 2a; social: *r* = 0.77; Fig. 2c; both *p* < *e*^*−10*^). For individual subjects, the topographies derived by TOPF (i.e., subject-specific PC1 loadings across ROIs), on average, achieved a moderate correspondence with those derived by GLM (motor: *r* = 0.50 ± 0.15, social: *r* = 0.64 ± 0.09; Supplementary Fig. 1a and b). Whereas TOPF conceptualises individual topographies as reflecting IEs of a shared response, thus ignoring the idiosyncratic response as captured by GLM, further results show that TOPF captured greater individual differences than GLM (Supplementary Fig. 1c).

**Fig. 2:**
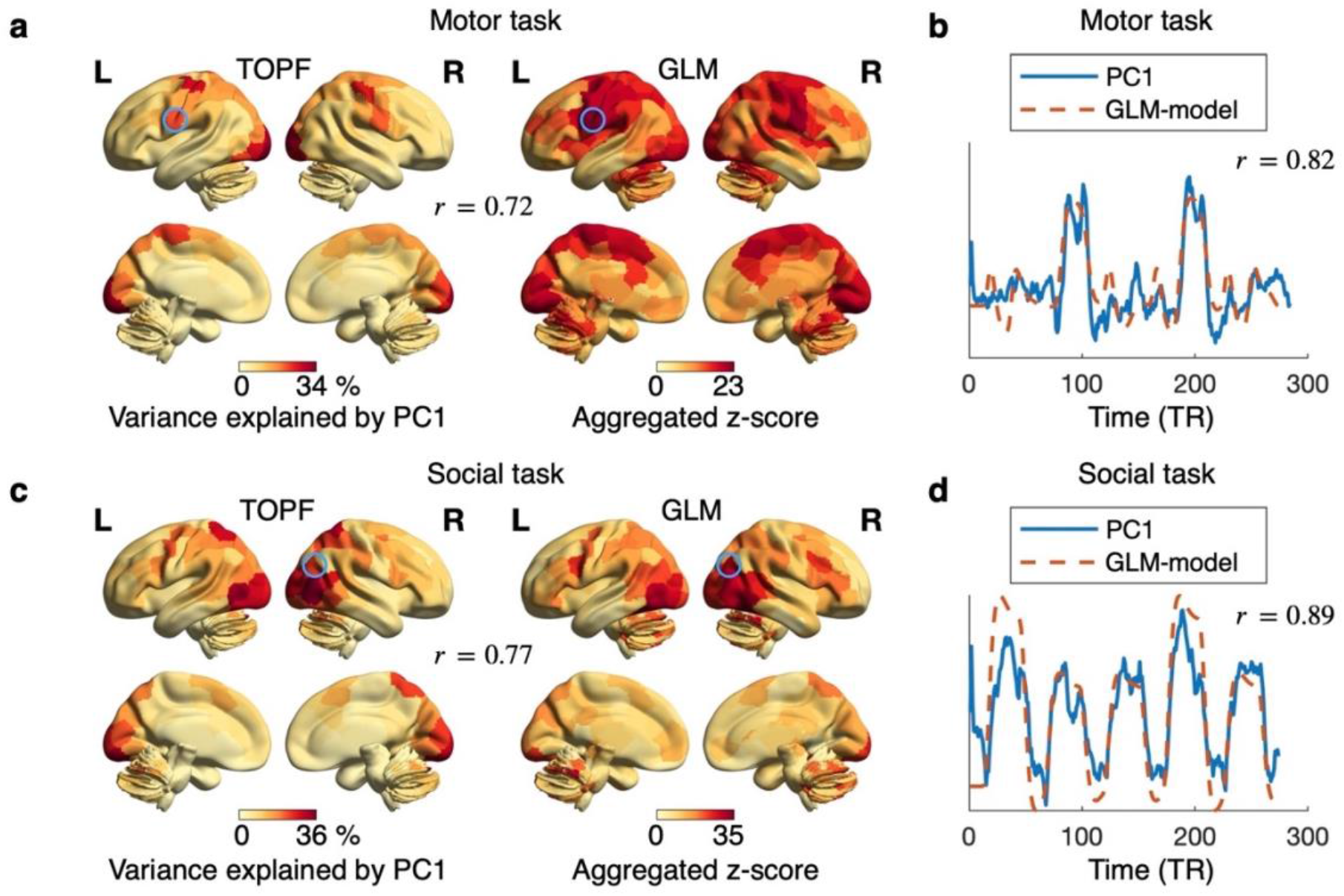
Validation of TOPF in capturing stimulus-evoked brain activity on task-based fMRI data. Correspondence (Pearson’s correlation coefficient, r) between TOPF-derived topographies and GLM-derived activation maps for the motor (**a**) and social (**c**) tasks separately at the group level. For TOPF, each value represents the amount of variance explained by the PC1 time series in the given ROI. For GLM, each value represents the activation strength (z-score) aggregated across all experimental conditions within each task (maximum of their absolute values) for the given ROI. The colour from yellow to red indicates the value from low to high. Correspondence (r) between the detected PC1 time series (blue) and the model used in GLM (red) for representative ROIs marked in circles in (a) and (c) for the motor (**b**) and social (**d**) tasks separately. For GLM, the model of each task is computed as the convolution of the HRF with the temporal structure of the given task aggregated over all experimental conditions. The brain maps are visualised using BrainNet Viewer [*68*].

The validity of the shared stimulus-evoked activity time series derived by TOPF was assessed for each task. Specifically, for each ROI, the detected PC1 time series was compared against a combination of models used in GLM, i.e., the timing of the relevant events convolved with a canonical hemodynamic response function (HRF) aggregated across experimental conditions. The stimulus-evoked activity time series derived by these two approaches showed high Pearson’s correlation coefficients in those strongly activated ROIs, such as the tongue movement-related left premotor cortex [*26*] for the motor task (*r* = 0.82; Fig. 2b) and the social cognition-related right temporoparietal junction (TPJ) [*27*] for the social task (*r* = 0.89; Fig. 2d; see Supplementary Fig. 2 for more results). Notably, as we expected, the correspondences between TOPF- and GLM-derived stimulus-evoked activity were higher when using PC1 than PC2 and PC3 for TOPF (Supplementary Fig. 3), demonstrating that PC1 is an appropriate choice for TOPF to capture task-evoked activity in this study.

### Evaluating the stability of TOPF on NV fMRI data

In TOPF, a stable identification of the shared response time series is essential for a meaningful characterisation of individual-specific topographies. To evaluate how the shared responses detected by TOPF change with sample composition and sample size, we used fMRI data of all available subjects from the HCP 7T subset (dataset2, *n* = 179; Supplementary Table 1) acquired while watching three different movie clips, namely “Two Men” (Movie1), “Welcome to Bridgeville” (Movie2), and “Pockets” (Movie3). For each ROI and movie clip separately, we measured the stability of the derived PC1 across 100 different subsamples over a range of sample sizes (*n*) from 10 to 90 by using Pearson’s correlation. To avoid possible bias from the family structure of this dataset [*21*], no subjects from the same family were included in the same subsample.

We quantified the overall stability as the mean stability across all ROIs (Fig. 3a). Given enough samples, the overall stability achieved a high value for all three movies with small variability across subsamples (at *n* = 90; Movie1: 0.90 ± 0.01; Movie2: 0.83 ± 0.02; Movie3: 0.84 ± 0.02). In particular, the stability of the PC1 time series achieved above 0.80 at *n* = 90 for most ROIs in the sensory, frontal and parietal cortices (Fig. 3b; Supplementary Fig. 4). We further confirmed the statistical significance of these PC1 time series by using permutation tests (Supplementary Fig. 5). For PC2 and PC3, their overall stability was much lower than that of PC1 (Supplementary Fig. 6), and their group-level topographies also showed distinct patterns from those of PC1 (Supplementary Fig. 7).

**Fig. 3:**
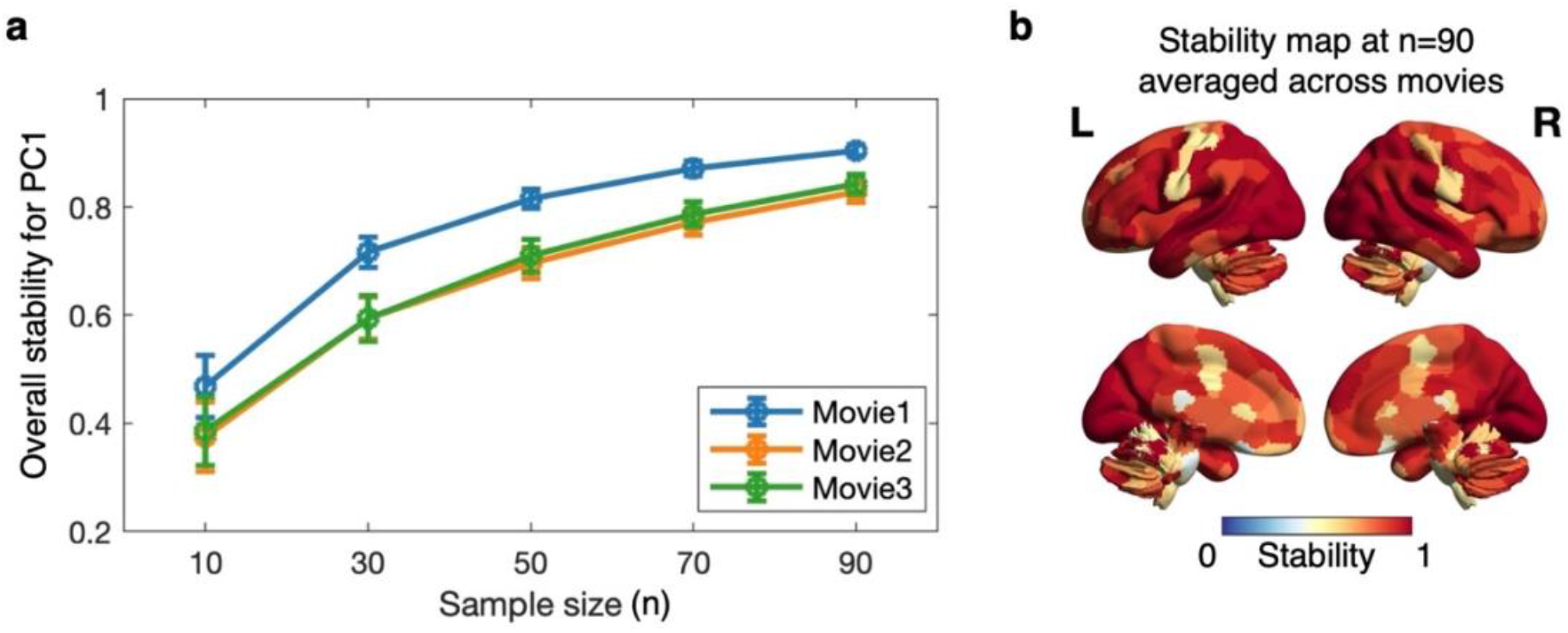
Stability of TOPF on NV fMRI data. **a**, Stability of the PC1 time series across subsamples over a range of sample sizes (*n*) from 10 to 90. The overall stability for PC1 is computed as the pairwise Pearson’s correlation coefficients averaged across all subsample pairs and then across all ROIs for each movie clip separately. The error bars depict the standard deviation across subsample pairs. **b**, Stability map of the PC1 time series across the whole brain at *n* = 90. For each ROI, the stability is averaged across all three movie clips. The colour from blue to red indicates the stability from low to high.

### Evaluating individual differences in individual-specific topographies on NV fMRI data

As TOPF reflects individual differences by IEs of shared responses rather than captures the idiosyncratic stimulus-evoked brain activity for each subject, it is necessary to evaluate whether such characterisation leaves enough room for individual variation. The individual-specific topographies derived by TOPF at *n* = 90 on the NV fMRI data stated above were used for this evaluation (see Fig. 4a for an illustrative example). For each pair of subjects, we computed their similarity in individual topographies by Pearson’s correlation within each subsample for each movie clip separately (Fig. 4b). On average, the between-subject similarity (*r*) achieved a relatively low value for all three movie clips (Movie1: *r* = 0.47 ± 0.14; Movie2: *r* = 0.41 ± 0.17; Movie3: *r* = 0.38 ± 0.18). Further results show that individual differences came from ROIs throughout the whole brain except the sensory cortex (Supplementary Fig. 8). These results demonstrate that these individual topographies captured substantial individual differences.

**Fig. 4:**
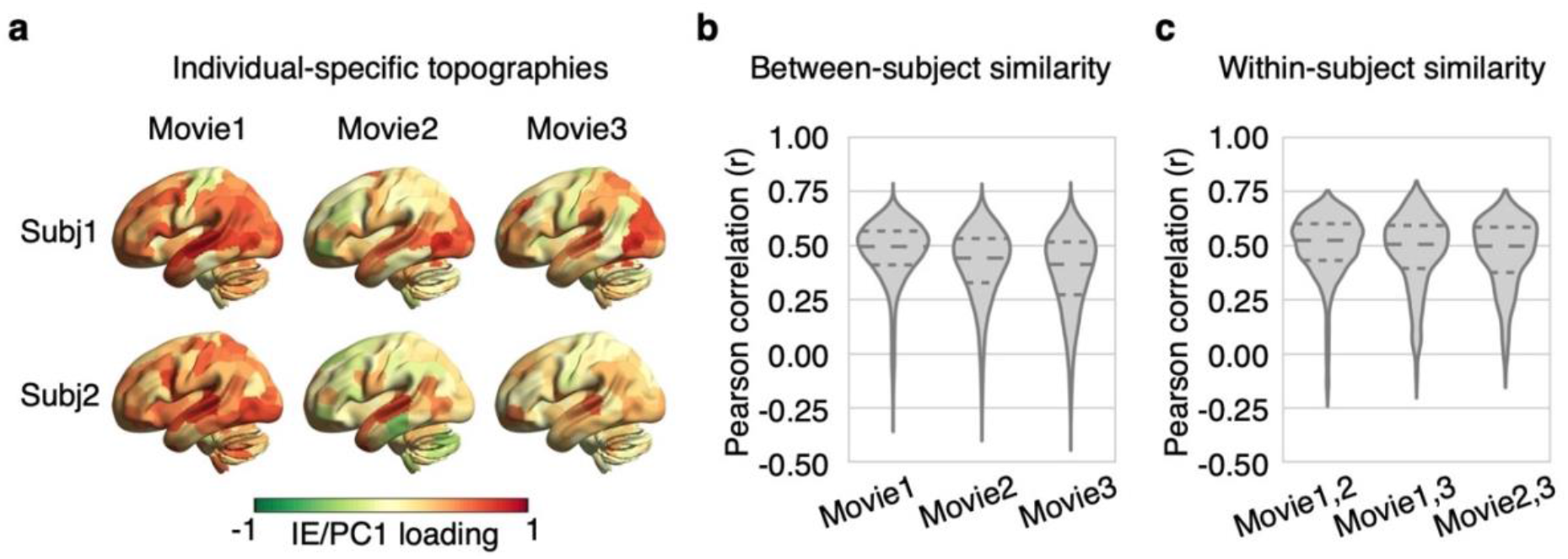
Individual differences in TOPF-derived individual-specific topographies on NV fMRI data. **a**, Individual-specific topographies of two representative subjects derived on a randomly selected subsample at *n* = 90 for each movie clip separately. The colour from green to red indicates the IE value (PC1 loading) from low to high. Each subject shows a unique spatial pattern of the IE values while watching the movie clips. **b**, Distribution of the between-subject similarity (Pearson’s correlation coefficient, *r*) in these individual-specific topographies over all pairs of subjects within each subsample (*n* = 90) for each movie clip separately. **c**, Distribution of the within-subject similarity (*r*) in the individual-specific topography over all subjects within each subsample (*n* = 90) for each pair of movies separately. The three dashed lines inside each violin plot from top to bottom denote the third quartile, median and first quartile of the corresponding distribution, respectively.

On the other hand, for each pair of movie clips, we computed the within-subject similarity for each subject on each subsample separately. The mean within-subject similarity (*r*) across subjects and subsamples achieved approximately *r* = 0.50 for all movie-clip pairs (Fig. 4c). In particular, for the ROIs in the sensory cortex [*28*], the pattern of the PC1 loadings over subjects remained relatively stable across movie clips (Spearman’s correlation *r* = 0.44 − 0.75; Supplementary Fig. 9). These results suggest that these individual topographies may reflect stable personal traits to a certain degree.

### Predicting individual phenotypes from individual-specific topographies

The second step in TOPF is to link the identified individual-specific topographies to individual phenotypes using a machine learning framework. A total of 8 phenotypes across cognition (fluid intelligence and working memory), personality (openness, agreeableness, conscientiousness, extraversion and neuroticism) and emotion (emotion recognition) were investigated (Supplementary Table 2). For prediction of each phenotype, we applied TOPF to fMRI data of 7 different paradigms from the same cohort of subjects (*n* = 179; dataset2) separately, including the three NV paradigms (Movie1-3) and four conventional tasks of similar scan durations, namely the motor, social, working memory, and language tasks (Supplementary Table 1). Prediction was performed by using a ridge linear regression model for its simplicity and robustness. The model was evaluated by a nested 10- fold cross-validation (CV) procedure with 10 repetitions. In each fold, the model was fitted on the training set by using the individual-specific topographies (i.e., the PC1 loadings of all ROIs) as features. The hyperparameter was optimised via an inner 5-fold CV. The fitted model was then tested by predicting the phenotypes of the test set. To avoid data leakage from the test set to the training set, features of the test set of each fold were derived based on the shared response learned on the training set, and subjects from the same family were ensured to stay either in the training or the test set. Prediction accuracy was assessed by the Pearson’s correlation coefficient (*r*) between predicted and observed scores over all subjects, with potential confounds (age, sex and head motion; Supplementary Table 3) being regressed out from both scores [*29*].

The prediction accuracy was significant for fluid intelligence, working memory, openness and emotion recognition (permutation-based *p* < 0.05, 5000 iterations; Fig. 5; see Supplementary Tables 4 and 5 for results of all phenotypes). Our subsequent analyses will thus mainly focus on these four phenotypes. Notably, the prediction performance varied substantially across fMRI paradigms. For example, working memory and openness were best predicted by Movie2 (*r* = 0.30±0.04) and Movie3 (*r* = 0.24±0.04), respectively, whereas fluid intelligence and emotion recognition were best predicted by the language task (*r* = 0.27±0.04) and social task (*r* = 0.23±0.02), respectively. Similar results were obtained before confound removal (Supplementary Table 6) and using the leave-one-out CV (Supplementary Table 7). Besides, for the NV fMRI data, overall, PC1-based topographies outperformed PC2- and PC3-based topographies for predicting the cognitive scores, and the combinations of these topographies slightly increased the prediction accuracies (Supplementary Fig. 10).

**Fig. 5:**
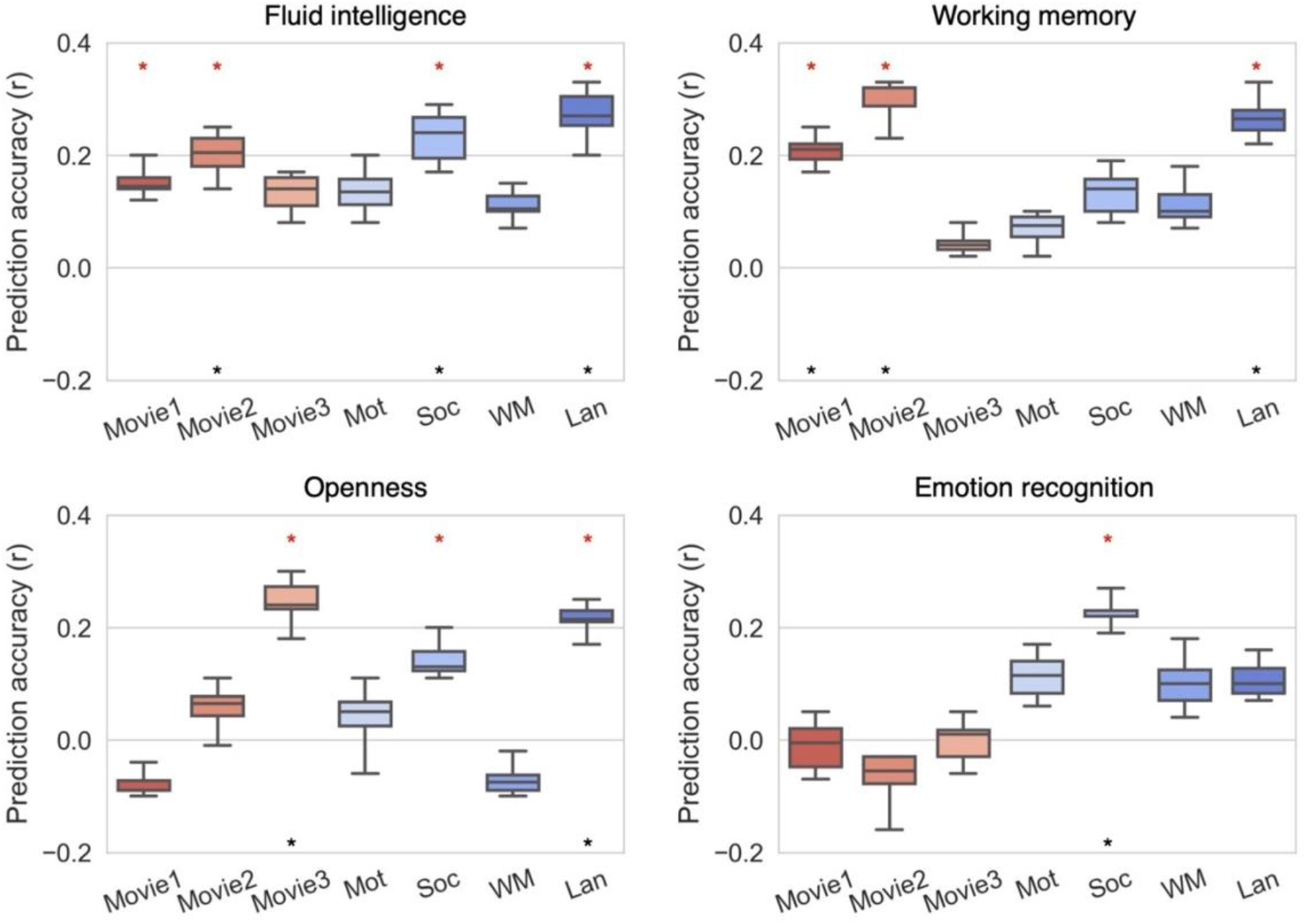
Prediction performance of TOPF evaluated in a 10-fold CV with 10 repetitions. Prediction accuracy is computed as the Pearson’s correlation coefficient (*r*) between predicted and observed scores over all subjects (*n* = 179) for each CV and fMRI paradigm-phenotype combination separately (box: middle bar, median; upper and lower bars, third and first quartiles; upper and lower whiskers: maximum and minimum). Sex, age and head motion are regressed out from both predicted and observed scores before computing the prediction accuracy. Significant predictions (*p* < 0.05, evaluated by permutation tests with 5000 iterations) before and after FDR correction are marked by red stars and black stars, respectively. NV and task paradigms are marked in red and blue, respectively. Only the phenotypes with significant predictions are shown here (see Supplementary Tables 4 and 5 for the complete results). Mot: motor task; Soc: social task; WM: working memory task; Lan: language task.

### Comparisons with functional connectivity-based prediction

Functional connectivity has so far been the most popular feature type for fMRI-based phenotype prediction. Here, we compared the prediction performance of TOPF with that of two commonly used functional connectivity-based feature sets, namely, the whole-brain connectome (WConn) and nodal connectivity strength (NConn). With the same procedure for performance evaluation being used, TOPF achieved better performance for predicting fluid intelligence and openness on most NV and task fMRI paradigms than WConn and NConn (Fig. 6). In particular, the highest accuracy was achieved by TOPF for all phenotypes. The performance of TOPF on NV and task data was also substantially better than that of the other two approaches when using RS data (Supplementary Fig. 11).

**Fig. 6:**
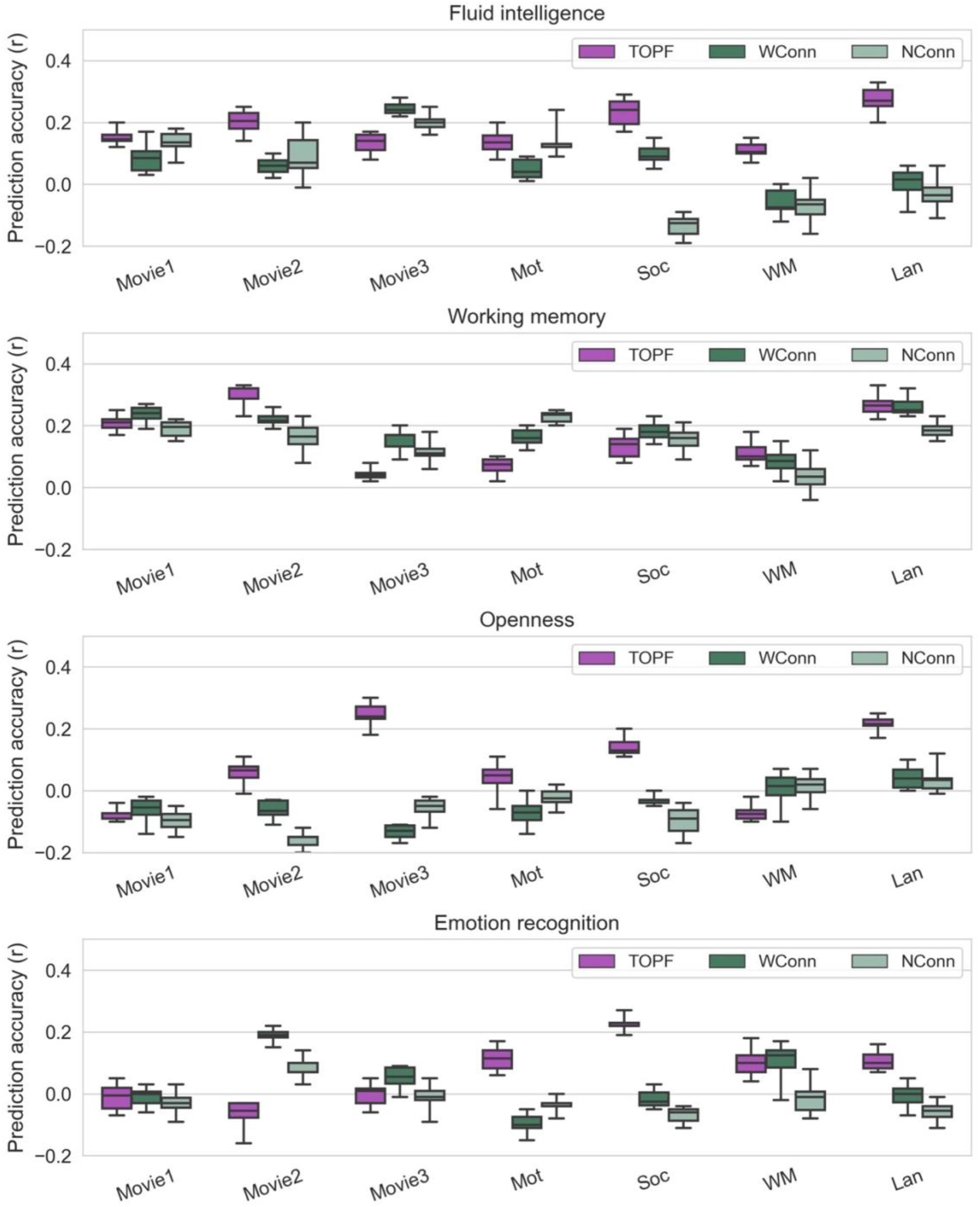
Comparisons between TOPF and connectivity-based prediction. For each phenotype, we show the distribution of prediction accuracy (*r*) over all 10 CV repetitions for each fMRI paradigm and each approach separately. The prediction accuracy is computed in the same way as computed in Fig. 5. Box plot: upper and lower whiskers: maximum and minimum; bars within each box from top to bottom: third quartile, median and first quartile. TOPF: PC1 loadings of all ROIs as features; WConn: whole-brain connectome as features; NConn: nodal connectivity strengths as features.

### Phenotype-related brain regions

To understand the neurobiological interpretation behind the prediction models obtained by TOPF, we identified the ROIs that were predictive of these phenotypes based on their permutation feature importance [*30*]. For each phenotype, we identified the predictive ROIs only for the fMRI paradigms in which the prediction accuracy was significant. For fluid intelligence, we identified a broad range of predictive ROIs, spreading across the frontal, parietal, temporal and cerebellar cortices [*31*] (Fig. 7a). For working memory, the identified ROIs were mainly located in the prefrontal, parietal, medial temporal and cerebellar cortices [*32*] (Fig. 7b). For openness, the identified ROIs were mainly located in the left frontal lobe and cerebellum [*33*] (Fig. 7c). For emotion recognition, the identified ROIs were mainly located in the right hemisphere, covering the frontal cortex, TPJ and subcortical regions [*34*] (Fig. 7d). Furthermore, different fMRI paradigms exhibited distinct spatial patterns of predictive ROIs for the same phenotype (Jaccard similarity: 0.02 to 0.10; Fig. 7e).

**Fig. 7:**
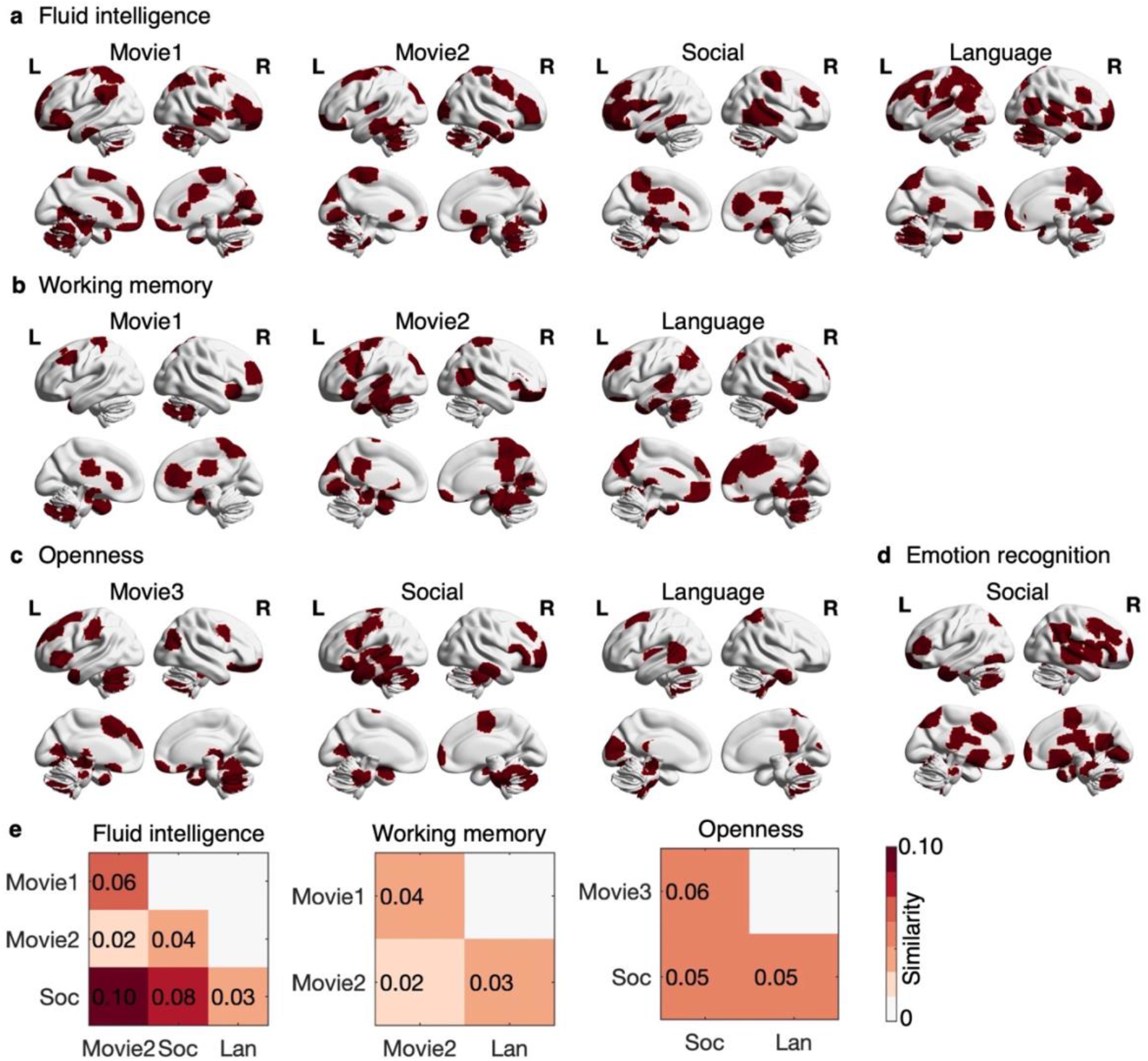
Predictive ROIs for each phenotype and fMRI paradigm with a significant prediction accuracy. **a-d**, The predictive ROIs (marked in red) are identified for each fMRI paradigm with a significant prediction accuracy (*p* < 0.05 in Fig. 5) for fluid intelligence (**a**), working memory (**b**), openness (**c**) and emotion recognition (**d**) separately. Only the ROIs for which the permutation feature importance over all CV folds and repetitions is significantly larger than zero (permutation-based, corrected *p* < 0.05, with 5000 permutations) are identified as predictive ROIs. **e**, The similarity in the spatial patterns of the predictive ROIs is assessed by the Jaccard similarity for each pair of fMRI paradigms within each phenotype separately. No result is plotted for emotion recognition, as only one fMRI paradigm (the social task) achieves a significant prediction accuracy.

## Discussion

The nascent technique, NV fMRI, shows great potential to advance our understanding of individual differences in complex brain functional processes under naturalistic conditions. To facilitate understanding individual differences in brain activity during NV, we proposed this novel, data-driven approach, TOPF, with unique features of NV fMRI data in mind. The TOPF approach allows us to characterise brain activity topographies evoked during NV in individuals and link them to individual phenotypes using machine learning techniques. TOPF effectively and stably identified the stimulus-evoked brain activity shared across subjects in a data-driven manner without requiring detailed stimulus descriptions. The individual-specific topographies conceptualised by TOPF provided a practical characterisation for individual activity patterns. These individual topographies predicted multiple individual phenotypes and outperformed the commonly used functional connectivity-based features. Last, by using TOPF, we were able to identify phenotype-related brain regions and provide neurobiologically meaningful interpretations.

While classic ICA and functional connectivity-based approaches are useful for studying individual differences in the interactions among brain regions, individual differences in brain activity directly driven by the stimulus, that is, the stimulus-evoked activity, might be of more interest for NV fMRI studies. On the other hand, the utility of GLM-based approaches to capture individual stimulus-evoked activity on NV fMRI data is limited by their requirement for accurate stimulus descriptions. Our TOPF approach avoids these issues by identifying stimulus-evoked activity in a data-driven way (Fig. 1). It detects a stimulus-evoked time course shared across subjects for each ROI separately and characterises individual activity patterns by the individual-specific expression (termed IE) of these shared responses across ROIs (termed individual-specific topography).

Similar to GLM, TOPF also defines a (few) consistent response time course(s) across subjects (TOPF: PC time courses; GLM: predefined regressors) and uses them as a common basis to estimate individual activity (TOPF: PC loadings; GLM: regression coefficients). However, in contrast to the model-driven manner used by GLM, TOPF can effectively and stably detect stimulus-evoked brain activity in a data-driven way without relying on stimulus descriptions (Figs. 2 and 3); hence, TOPF is highly suitable for dealing with fMRI data from complex tasks (e.g., movie watching). Notably, whereas GLM picks up individual idiosyncrasies by using predefined models to fit individual fMRI signals, TOPF uses IEs of a shared response to reflect individual variation. Even so, TOPF still leaves enough room for preserving individual differences (Supplementary Fig. 1). Moreover, the data-driven nature may allow TOPF to capture unique individual differences that are not induced by explicitly predefined task designs [*35*] and thus may not be detected by GLM.

TOPF is also closely related to several existing data-driven approaches for detecting stimulus-evoked activity. A common rationale behind these data-driven approaches is that any temporal variance shared across subjects can only originate in the processing of the same stimulus. For example, the commonly used intersubject correlation (ISC) approach [*5*] often computes the pairwise or leave-one-out (LOO) correlation (i.e., correlation between the fMRI time series of a subject and the time series averaged across the other subjects) to reflect the similarity in brain activity between subjects [*6, 10*]. Although similar, the PC1 loadings used in TOPF are more computationally efficient than the LOO correlations [*22*], thus allowing for more efficient integrations with machine learning models, especially when the number of subjects/ROIs is large. More importantly, TOPF aims to use an “absolute” expression of a group-level basis to reflect individual activity instead of providing a relative measure. While SRM [*19*] and hyperalignment [*20*] approaches adopt similar logic, they often focus on more fine-grained individual functional topographies within certain brain regions rather than the whole-brain topographies. Tensor-ICA [*11, 36*], on the other hand, identifies a set of common stimulus-evoked spatial components throughout the whole brain and associated time courses, with individual differences being characterised for each component. By contrast, TOPF characterises individual variation by ROI, delineating individual topographies across the whole brain at a resolution suitable for machine learning-based predictions.

It is important to note that TOPF characterises individual variation by IEs of stimulus-evoked responses that are shared across subjects rather than capturing individual idiosyncratic stimulus- evoked responses. Although the latter might be conceptually more desirable, we show that the IEs also captured considerable and meaningful individual differences (Fig. 4). These IEs were more variable across subjects in frontoparietal, limbic, subcortical and default mode network regions than in sensory regions (Supplementary Fig. 8). These results partially align with previous studies observing higher intersubject variability in functional connectivity patterns of brain regions associated with cognitive control in both RS [*37*–*39*] and NV settings [*40*]. Moreover, for the sensory regions, although the intersubject variability was relatively small, patterns of the IEs across subjects were stable across movie clips and thus less sensitive to the specific movie content (Supplementary Fig. 9). This is in line with recent studies showing that the way the brain processes complex sensory and social information during NV may be an intrinsic characteristic of individuals [*8, 28*] and associated with brain baseline functional organisation [*41*]. We note that further work is needed to assess whether our findings here are generalisable across different movie types.

Using machine learning predictive models to promote understanding of brain-behaviour relationships and identification of biomarkers of brain diseases has recently gained considerable popularity [*15*– *17*]. While many studies have successfully predicted individual phenotypes from RS, task fMRI and other neuroimaging data, NV settings are expected to further promote individual phenotype predictions due to their favourable properties (e.g., enhancing individual differences) [*40*]. Indeed, recent studies have found that various phenotypes and diseases are related to individual variations in brain activity during NV [*6, 11, 12*]. With TOPF, we extend these previous findings by showing that individual activity during NV can predict a variety of individual phenotypes using machine learning frameworks on novel subjects (Fig. 5). In addition, TOPF also performed well on phenotype-relevant conventional tasks [*42*], further highlighting the utility of TOPF for phenotype prediction.

Several recent studies have predicted individual phenotypes on NV fMRI data from individuals’ functional connectivity profiles [*43*]. In our present study, we also compared the performance of TOPF to the prediction performance of two commonly used functional connectivity-based feature sets (Fig. 6). Compared to using whole-brain connectomes as features, TOPF achieved better performance for openness and similar performance for cognitive scores by using a remarkably smaller number of features (i.e., *n* ROIs instead of *n*(*n* − 1)/2 connections). This greatly eases the problem of overfitting and reduces the number of observations (i.e., subjects) needed for a meaningful prediction, alleviating the critical problem of relatively small sample sizes of the current public NV fMRI datasets. In addition, TOPF achieved substantially better performance than the nodal connectivity strength-based prediction on both NV and task-based fMRI data, although these two approaches have the same number of features. On the other hand, the feature type achieving the best performance was highly variable across phenotypes and tasks/movies, indicating that these different types of features may reflect unique and complementary aspects of brain functions [*44*]. These findings may shed light on the open question of which types of features best suit phenotype prediction on NV fMRI data.

In addition to accuracy, a useful prediction model is also expected to have good interpretability, especially in neuroscience studies. Our results demonstrate that TOPF holds great potential for facilitating our understanding of the neurobiological bases of behaviour by localising phenotype-related brain regions (Fig. 7). For example, the ROIs identified for fluid intelligence and working memory were mainly located in brain regions supporting cognitive functions, e.g., frontal and parietal cortices [*45*], showing overlaps with findings of previous meta-analytic studies [*32*]. The association between the left frontal lobe and the openness personality trait has been reported by previous studies using conventional task fMRI data and brain structural data [*46, 47*]. The TPJ identified for emotion recognition has been recognised as a key region for social and emotion processing [*27*]. Besides, the patterns of predictive ROIs identified for TOPF varied largely across fMRI paradigms, indicating that different tasks and movies may elicit unique neural processes related to the same behaviour or phenotype [*48*]. We expect that future studies on the important question of whether and how brain function under naturalistic conditions differs from that measured in strictly controlled task paradigms can be facilitated by using our approach.

It is noteworthy that, first, while our experimental results here show that PC1 outperformed PC2 and PC3 in phenotype prediction in most cases (Supplementary Fig. 10), this does not indicate that the residual fMRI signals after regressing out PC1 have limited behaviourally relevant information; rather, it highlights the need for more advanced approaches to characterise meaningful individual differences therein. The other PCs may also be useful for future studies, for instance, when subjects are from different age groups showing distinct shared response patterns across groups [*22, 37*]. Second, although TOPF assumes a (few) shared response(s) for each ROI, this may not be the case for certain ROIs (e.g., those that are not responsive to the movie stimulus). Incorporating feature selection techniques or a priori knowledge into TOPF may further optimise the performance of TOPF for phenotype prediction [*49*]. Finally, whole-brain parcellations play an important role in TOPF because they can reduce the spatial dimensionality of fMRI data in a biologically meaningful way [*14*] and ease the computational load for subsequent analyses. As the best-suited parcellations may vary across different brain states and for answering different research questions [*5*0], it will be useful to evaluate the influence of different parcellation schemes on the performance of TOPF in the future.

In sum, the TOPF approach presented here provides a data-driven, machine learning-based tool for studying individual differences in evoked brain activity. Essentially, TOPF characterises individual multivariate evoked activity patterns and highlights their value for understanding brain-behaviour relationships on NV fMRI data. Although in this study TOPF was only tested on healthy participants, this principled and flexible approach should be readily adapted for clinical applications. We envision that TOPF will provide a powerful tool for not only predicting symptom severity or clinical outcomes but also identifying potential biomarkers in clinical neuroscience studies.

## Materials and Methods

### Datasets

All participants used in this study were from the HCP S1200 release [*21*]. Specifically, the HCP “100 Unrelated Subjects” subset (dataset1) was used for the validation of TOPF in capturing stimulus-evoked brain activity, and the HCP 7T subset (dataset2) was mainly used for the evaluation of TOPF in phenotype prediction. Details of the fMRI and behavioural data included in the two datasets are described below (see Supplementary Tables 1 and 2 for an overview). Informed consent was obtained from all participants [*21*].

Dataset1 was from the HCP “100 Unrelated Subjects” subset (n = 100; age range: 22-36 years; mean age = 29.11±3.68 years; 54 females/46 males) of the full HCP dataset. The conventional task-based fMRI data in this dataset were used to validate the ability of TOPF to capture stimulus-evoked brain activity due to the availability of the temporal structure of these task designs. This dataset was used because the subjects in this dataset were unrelated, which straightforwardly avoided the possible bias from the genetic relatedness in the full HCP dataset. Each subject was required to complete seven different tasks during the scan in two fMRI sessions, and each task was performed in two runs with different phase-encoding directions. Among these tasks, we chose the motor task and the social cognition (social) task as two representative tasks because they tapped into two different levels of the brain functional hierarchy: whereas the motor task mainly involved simple movements of body parts, the social task involved one higher-order cognitive function of the brain, i.e., the theory of mind ability. Protocol details about these tasks were described by previous studies [*24*]. For both tasks, we limited our analysis to the data from the first run (with right-to-left phase encoding), where the scan duration was 3’34” (284 TRs) for the motor task and 3’27” (274 TRs) for the social task. fMRI images were acquired using a 3T Siemens scanner (TR = 720 ms, TE = 33.1 ms, resolution = 2.0 mm^3^). Details of the acquisition parameters can be found elsewhere [*21*].

Dataset2 was from the HCP 7T subset (n = 184; age range: 22-36 years; mean age = 29.43±3.35 years; 112 females/72 males), which included all available NV fMRI data in the full HCP dataset. Different from dataset1, subjects in dataset2 contained twins and siblings from 93 unique families. Each participant underwent four NV fMRI runs in two sessions. In each run, subjects were presented with a sequence of different short movie clips (4-5 clips) interleaved with 20 s rest blocks, resulting in a total duration of approximately 15 mins for each run. All NV fMRI images were acquired using a 7T Siemens scanner (TR = 1000 ms, TE = 22.2 ms, resolution = 1.6 mm^3^). Detailed descriptions of the acquisition parameters, imaging protocols and experimental stimuli can be found elsewhere [*21, 43*]. In this study, we limited our analysis to the NV fMRI data from the first run because the movie stimuli used in the other three runs had slight differences in timing across subjects, which may potentially affect the results. fMRI data during watching of the last two movie clips in the first run were also excluded due to the short scan duration (less than 2 mins). As a result, we only included fMRI data acquired while watching the first three movie clips of the first run, i.e., “Two Men” (Movie1), “Welcome to Bridgeville” (Movie2), and “Pockets” (Movie3), for which the scan duration was 4’04” (244 TRs), 03’41” (221 TRs) and 03’08” (189 TRs), respectively.

Apart from the NV fMRI data, we also included task-based fMRI data from the same subjects in dataset2 to evaluate the performance of TOPF in phenotype prediction. Consistent with dataset1, these task-based fMRI images were also acquired using a 3T Siemens scanner with the same experimental settings, imaging protocols and parameters (TR = 720 ms, TE = 33.1 ms, resolution = 2.0 mm^3^). For a more straightforward comparison between NV and task-based paradigms, we only included the tasks for which the scan durations were similar to those of the NV paradigms (3-5 mins), which were the motor (3’34”), social (3’27”), working memory (5’01”) and language tasks (3’57”). Similarly, only the data from the first run (with right-to-left phase encoding) were used here.

In addition, the RS fMRI data of all the subjects in dataset2 were also included in our analysis for comparison. These RS fMRI data were acquired before the NV fMRI run we used in this study (described above) in the same session using a 7T Siemens scanner with the same imaging protocols and parameters. The scan duration of the RS fMRI data was 15 mins (900 TRs). We note that as there is no external stimulus used in RS settings, our TOPF method is not applicable to RS fMRI data. Instead, two functional connectivity-based approaches were applied to the RS fMRI data and compared to the TOPF method in terms of the prediction performance (see the “Comparisons with functional connectivity-based features for phenotype prediction” section below for more details).

The behavioural phenotypes of the subjects in dataset2 studied here included 8 different phenotypes covering fluid intelligence, working memory, five dimensions of personality and emotion recognition (Supplementary Table 2). Detailed descriptions of the tests to measure these phenotypes can be found elsewhere [*24*]. Note that five subjects were excluded due to the lack of the complete fMRI and behavioural data mentioned above, resulting in a total of 179 subjects (108 females/71 males) for dataset2.

### fMRI data preprocessing

All 3T task-based fMRI images were preprocessed with the HCP minimal preprocessing pipeline [*51*], which includes gradient unwarping, motion correction, spatial normalisation to the Montreal Neurological Institute (MNI) space and intensity normalisation. We further preprocessed these images by regressing out Friston’s 24 head motion parameters [*52*], as well as the mean time series of white matter and cerebrospinal fluid and the linear trend, using the Data Processing and Analysis for Brain Imaging (DPABI) toolbox [*53*] (http://rfmri.org/dpabi) in Matlab R2019a. All 7T NV and RS fMRI images were preprocessed with the standard HCP pipelines [*51*], including correction for distortion and motion, registration to the MNI space, high-pass filtering, removal of 24 motion parameters and FIX-denoising [*54*]. The first 10 volumes were discarded to correct for hemodynamic delays of the NV fMRI data for each movie clip separately. For all the NV, task-based and RS fMRI images, the whole brain was divided into 268 ROIs using a functionally defined parcellation that has been widely applied to the HCP dataset [*23*]. The parcellation was resampled to match the spatial resolutions of the corresponding fMRI images. Within each ROI, the mean time series over all voxels was extracted and z-score normalised (i.e., zero-mean with unit-variance) for each subject. Head motion was measured by the relative root-mean-square framewise displacement (FD) for each subject and each fMRI paradigm. The FD values of all subjects from all these fMRI paradigms were less than 0.5 mm, and thus no subject was further excluded from the analysis.

### Formulation of TOPF: identifying individual-specific topographies

The first step of TOPF is to delineate individual stimulus-evoked brain activity patterns across ROIs from NV fMRI data in a data-driven manner. To do this, we modified a commonly used model of NV fMRI signals. Specifically, it is often assumed that, in each brain region, for a given subject *i*, the observed NV fMRI time series *x*_*i*_ consists of three time series components [*10*]:

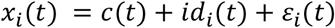

where *c* is a stimulus-evoked component that is shared across all subjects (denoted as shared response), *id*_*i*_ is also stimulus-evoked but unique to each individual (denoted as idiosyncratic response), *ε*_*i*_ is the residual representing the other signal components, and *t* is a specific time point. Inspired by recent studies [*6, 22*], we modified the above formulation as follows:

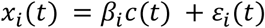

such that the idiosyncratic information related to the stimulus-evoked brain activity is summarised as an individual-specific scalar, *β*_*i*_. This value *β*_*i*_ represents the individual-specific expression level of the shared response time series (*c*) for subject *i*, which we refer to as IE. By using *β*_*i*_*c*(*t*) instead of *c*(*t*) + *id*_*i*_(*t*), we defined a common basis (*c*) to represent the individual-specific information in the stimulus-evoked brain activity as a single value *β*_*i*_.

To identify the shared response time series *c* and the IE value *β*_*i*_, first, for each ROI, we constructed a data matrix of preprocessed fMRI time series across subjects, *X* = [*x*_1_, *x*_2,_…, *x*_*n*_] ∈ *R*^*d*×*n*^, where *n* is the number of subjects, *d* is the number of time points, and *x*_*i*_ is the z-score normalised time series for subject *i*. Next, a PCA decomposition was performed on *X* with the subjects being treated as variables. The resulting PC time series were used as an estimate of the shared response time series *c*, and the subject-wise loadings onto the PCs were used to reflect the IE values (i.e., *β*_*i*_) of *c*. This procedure was repeated for each ROI separately. Finally, for each subject, we collected the PC loadings of that subject across all ROIs to reflect a brain activity pattern specific to that subject, a construct we refer to as individual-specific topography. The PCA decomposition was implemented in Python using the “*sklearn*.*decomposition*.PCA” function in the scikit-learn package [*55*] (https://scikit-learn.org/stable/). Because PC1 explains the largest amount of variance shared across subjects, in this study, we mainly used PC1 to reflect the shared response time series and operationalised the IE values as PC1 loadings. In the Supplementary Information, we further analysed the results for PC2 and PC3.

### Validation of TOPF

To assess the validity of TOPF in capturing stimulus-evoked brain activity, we used the task-based fMRI data in dataset1. First, for each task, we compared the group-level topography derived by TOPF with the group-level task-evoked activation (z-score) map derived by GLM by using Pearson’s correlation. The TOPF-derived topography was generated by computing the amount of variance explained by PC1 for each ROI separately and collecting these values across all ROIs. The GLM-derived activation maps were provided by the HCP and downloaded from NeuroVault [*56*] (https://neurovault.org). We expected the variance explained by PC1 to be an effective measure of the group-level task-evoked activation, under the assumption that a stronger group-level activation is associated with higher similarity in brain activity across subjects [*5*]. This assumption was supported by the high correspondences between the TOPF-identified topographies and the GLM-derived activation maps (Fig. 2a and c). The correspondences between the TOPF-derived individual-specific topographies (i.e., PC1 loadings across all ROIs) and the individual-level GLM-derived activation maps (downloaded from the HCP) were also assessed by using Pearson’s correlation for each subject separately (Supplementary Fig. 1a and b). We note that all these GLM-derived activation maps were initially generated for each experimental condition separately (e.g., left-hand movement or tongue movement), in contrast to the TOPF-identified topographies, which reflected the overall brain activity across all experimental conditions. Therefore, we aggregated the GLM-derived activation maps across all experimental conditions within each task for a more straightforward comparison, where the maximum of the absolute activation values was computed for each voxel in FSL [*57*] (http://fsl.fmrib.ox.ac.uk/fsl) and then averaged over all voxels within each ROI. The amount of individual differences captured by these individual-level topographies/maps was measured by between-subject correlations and compared between TOPF and GLM (Supplementary Fig. 1c).

Second, we compared the detected PC1 time series with the models used in GLM for each individual ROI separately. Specifically, for each experimental condition *i*, we modelled a canonical response (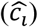) by convolving the timing of the relevant events with the canonical HRF as implemented by the “*spm_hrf*.*m*” function of the Statistical Parametric Mapping (SPM) software [*58*] with default settings in Matlab R2019a. For each ROI, we then combined these canonical responses across all experimental conditions with the GLM-derived activations of the given ROI in these conditions as weights:

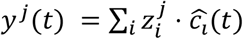

where *y*^*j*^(*t*) denotes the combined model of ROI *j* at time point *t* and 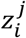 denotes the GLM-derived activation of ROI *j* in task condition *i*. The Pearson’s correlation coefficient between the combined model *y*^*j*^ and the PC1 time series was computed for each ROI in each task (Fig. 2b and d; Supplementary Fig. 2). The above analyses were also conducted for PC2 and PC3, for which the correspondences with the GLM-derived results were much lower than those of PC1 (Supplementary Fig. 3).

### Analysis of the stability of TOPF

The stability of the TOPF-derived shared response time series across different subsamples was evaluated on the NV fMRI data in dataset2. This is necessary because the individual-specific topographies are essentially determined by the shared response time series, and the latter may change across different subsamples due to the data-driven nature of TOPF. For each movie clip, we varied the sample size (number of subjects, *n*) of the input fMRI data matrix from 10 to 90 with a step size of 20 and created 100 different subsamples for each sample size. Only unrelated subjects were included in the same subsample to avoid possible bias from the family structure of the dataset. The stability was first evaluated for each ROI separately at each value of *n* by computing the mean Pearson’s correlation coefficient of the derived PC1 time series across all pairs of subsamples (Supplementary Fig. 4). Given that the derived PC1s may be sign-flipped for some subsamples, we used the absolute correlation value to reflect the similarity of any two PC time series. The stability was then averaged over all ROIs to reflect the overall stability across the whole brain (Fig. 3). As the overall stability of PC1 was substantially higher than that of PC2 and PC3 (Supplementary Fig. 6), we mainly focused on the results of PC1 for the following analyses in the main text.

### Analysis of the statistical significance of TOPF-derived shared responses

The statistical significance of the PC1 time series derived by TOPF was evaluated by using permutation tests on the subsamples (as described above) of the NV fMRI data at *n* = 90. This analysis determined whether the amount of variance explained by each PC1 time series exceeded the chance level. For each ROI on each subsample of each movie clip, we permuted the input fMRI data matrix *X* by applying circular shifting. The fMRI time series of each subject was shifted with a random time interval so that the fMRI time series across subjects were mismatched in time while the autocorrelation structure of each fMRI time series was preserved [*59*]. We next computed the variance explained by the PC1 time series of this permuted data matrix. This procedure was repeated 10000 times with different permutations of *X*, and the variance explained by PC1 was collected across all iterations to create a null distribution. A p-value was then determined as the proportion of iterations on which the variance explained by PC1 was larger than that derived on the non-permuted data and Bonferroni-corrected (Supplementary Fig. 5). The whole-brain topographies of the variance explained by PC1 were compared with those of the variance explained by PC2 and PC3 by using Pearson’s correlation (Supplementary Fig. 7).

### Evaluation of individual differences in individual-specific topographies

The ability of TOPF to capture individual differences was assessed by the between-subject similarity of individual-specific topographies (i.e., PC1 loadings across ROIs). The topography was derived by TOPF for each subject on each subsample (as described previously) of the NV fMRI data at *n* = 90. The between-subject similarity in these topographies was computed by using Pearson’s correlation for each pair of subjects within each subsample for each movie clip separately (Fig. 4b). For each individual ROI, intersubject variability in brain activity was quantified as the standard deviation of PC1 weights (Supplementary Fig. 8). Note that here we used PC1 weights for a fair comparison across different ROIs, which are equivalent to the PC1 loadings normalised by the square root of the eigenvalue of the given PC1. Likewise, the within-subject similarity was computed by the Pearson’s correlation coefficient of the topographies between each pair of movie clips for each subject within each subsample separately (Fig. 4c). The cross-movie stability of the pattern of the PC1 loadings across all subjects was computed by using Spearman’s correlation for each ROI and each pair of movie clips on each subsample separately (Supplementary Fig. 9).

### Phenotype prediction: inference of brain-behaviour relationships

The second step of TOPF is to investigate the behavioural relevance of the identified individual-specific topographies under a machine learning-based predictive framework. Seven fMRI paradigms from the NV and task-based fMRI data in dataset2 were used to predict eight different phenotypes separately to evaluate the predictive framework. In this study, we used a ridge linear regression model for the prediction. We chose this particular model because it is simple and has been shown to achieve robust performance in various neuroimaging studies [*60, 61*]. We note that other prediction models could also be used within TOPF in general.

Prediction models were trained in a 10-fold cross-validation (CV) scheme. Specifically, all subjects (*n* = 179) were randomly divided into 10 groups of roughly the same size. To account for the family structure of dataset2, subjects from the same family were ensured to be all included in either the training set or the test set for each fold. In each fold, one group was held out for testing, with subjects from the other groups being used for training. Each group was used for testing once and only once. In each fold, the training set was used to train a prediction model, where for each subject all IE values (PC1 loadings) of the individual-specific topography were used as features, resulting in a 1 × 2*6*8 feature vector. Each feature was then z-score normalised. To avoid information leakage from the test to the training set, individual-specific topographies of test subjects were derived based on the shared response time series learned on the training set. Specifically, for each ROI of each test subject, the IE value was computed as the Pearson’s correlation coefficient between that subject’s fMRI time series and the PC1 time series learned on the training set of the given ROI. We note that on the training set the PC loading (i.e., the feature) of each subject is mathematically equivalent to the Pearson’s correlation coefficient between the time series of that subject and the corresponding PC time series [*62*]. This whole procedure ensured that test subjects were totally separated from training subjects during both the feature extraction and phenotype prediction phases. Besides, for each training set, the optimal regularisation parameter *λ* of ridge regression was determined by an inner 5-fold CV from 12 values [2^−5^, 2^−4^, …, 2^*6*^]. The whole pipeline was implemented in Python using the scikit-learn and julearn packages (https://juaml.github.io/julearn/main/index.html). For each fMRI paradigm- phenotype combination, the above procedure was repeated 10 times with different groupings of subjects.

For each computation, prediction accuracy was quantified as the Pearson’s correlation coefficient between the predicted and the observed phenotypic scores across all subjects and all 10 folds. The accuracy was then averaged over the 10 repetitions. To control for the influence of head motion, age and sex [*63*], we regressed out these confounds from both the observed and predicted scores and computed the correlation of the two residuals as accuracy [*29*]. Permutation tests with 5000 iterations were used to assess whether the obtained prediction accuracy was significantly higher than the chance level. For each iteration, the predicted scores were randomly shuffled across subjects with the original orders of the confounds and observed scores kept. In this way, the relationship between the predicted and observed scores was destroyed while the relationship between the observed scores and confounds was preserved [*29*]. The correlation between the residuals of the permuted scores and observed scores after regressing out the confounds was computed for each iteration and collected across all iterations to construct the null distribution. A p-value was then obtained by computing the proportion of iterations for which the resulting prediction accuracy was higher than that on the non-permuted data (Supplementary Tables 4 and 5). The prediction accuracy of TOPF (Fig. 5) was slightly lower than that evaluated without confound removal (Supplementary Table 6) and highly similar to that derived using leave-one-out CV (Supplementary Table 7). For the NV fMRI paradigms, the performance of TOPF was further compared to that derived by using PC2 loadings, PC3 loadings, the concatenation of PC1 and PC2 loadings (PC1-2), and the concatenation of PC1, 2 and 3 loadings (PC1-3) as features separately (Supplementary Fig. 10).

### Comparisons with functional connectivity-based features for phenotype prediction

As functional connectivity profiles have been widely used for phenotype prediction on fMRI data, we adopted two commonly used functional connectivity-based features for prediction, i.e., the whole-brain connectome (WConn) and nodal connectivity strength (NConn), and compared their performance with TOPF. Apart from the seven fMRI paradigms (three NV paradigms and four tasks), we further included RS fMRI data of all the subjects in dataset2 (described above). For each subject and each fMRI paradigm, we computed the functional connectivity as the Pearson’s correlation coefficient of the fMRI time series for each pair of ROIs separately, resulting in a total of eight 2*6*8 × 2*6*8 connectivity matrices for each subject. Each correlation coefficient was then Fisher z-transformed. For WConn, we extracted the upper triangle of the connectivity matrix without the diagonal as features, yielding a total of 35778 features (connections) for each subject and each fMRI paradigm. For NConn, we computed the node strength for each ROI separately, which is defined as the sum of the absolute values of the connectivities between the given ROI and all the other ROIs, resulting in a total of 268 features (ROIs) for each subject and each fMRI paradigm. The WConn and NConn features were then used to predict fluid intelligence, working memory, openness and emotion recognition on each fMRI paradigm separately by repeating the same prediction pipeline as used in TOPF. Consistent with TOPF, the prediction accuracies were also computed by the Pearson’s correlation coefficients between the predicted and observed scores after controlling for the three confounding variables (Fig. 6; Supplementary Fig. 11).

### Identification of phenotype-related brain regions

Permutation feature importance [*30*] was computed for each ROI in each fMRI paradigm-phenotype combination with a significant prediction accuracy (*p* < 0.05 in Fig. 5) separately. Specifically, for each successful combination, we collected the models fitted on the training sets across all CV folds and all repetitions, resulting in a total of 100 models. For each ROI and each model, we randomly shuffled the observations of the given ROI in the test set corresponding to the model and recomputed the prediction accuracy on the permuted test set. After repeating the procedure 100 times, the importance of each ROI in the given model was then quantified as the decrease in prediction accuracy (i.e., the difference between the true accuracy and the accuracy derived on the permuted test set), averaged across the 100 iterations. This analysis was implemented in Python using the “*permutation_importance*” function in the scikit-learn package. Finally, for each successful fMRI paradigm-phenotype combination, a right-tailed, one-sample permutation test [*65*] with 5000 iterations was used to identify the ROIs for which the importance values over all the models were significantly larger than zero (corrected *p* < 0.05; Fig. 7a-d). The correction for multiple comparisons was performed against a null distribution of the maximum value across all ROIs in each iteration [*66*]. We note that in this study we used the permutation feature importance instead of the feature weights (i.e., regression coefficients) to identify the predictive features because the latter are known to have certain limitations in their interpretability [*61, 67*]. The similarity of the spatial patterns of the identified predictive ROIs between fMRI paradigms was computed by the Jaccard similarity index for each pair of fMRI paradigms with significant predictions for each phenotype separately (Fig. 7e).

## Data availability

The HCP data used in this study (preprocessed task-based and NV fMRI data, behavioural data, and GLM-derived activation maps for each single subject) are publicly available from the HCP website (https://db.humanconnectome.org/). Family structure information can be accessed after approval of the HCP Restricted Data Use Terms.

## Code availability

The code to implement TOPF and subsequent analyses will be made available upon publication of the manuscript at https://github.com/xuanli-ac/TOPF.

## Acknowledgements

This work was supported by the European Union’s Horizon 2020 Research and Innovation Programme under grant agreement no. 945539 (HBP SGA3), and the Deutsche Forschungsgemeinschaft (491111487). Data were provided in part by the Human Connectome Project, WU-Minn Consortium (Principal Investigators: David Van Essen and Kamil Ugurbil; 1U54MH091657) funded by the 16 NIH Institutes and Centers that support the NIH Blueprint for Neuroscience Research; and by the McDonnell Center for Systems Neuroscience at Washington University.

## Author contributions

X.L., S.W, and S.B.E. conceptualised the study; X.L. performed the analysis with support from P.F. and K.R.P.; S.W., P.F., S.B.E., and K.R.P. contributed to result interpretation; X.L. drafted the manuscript, with contribution from S.W. and P.F. and comments from S.B.E. and K.R.P.

