## Supplementary Information for "A topography-based predictive framework for naturalistic viewing fMRI"

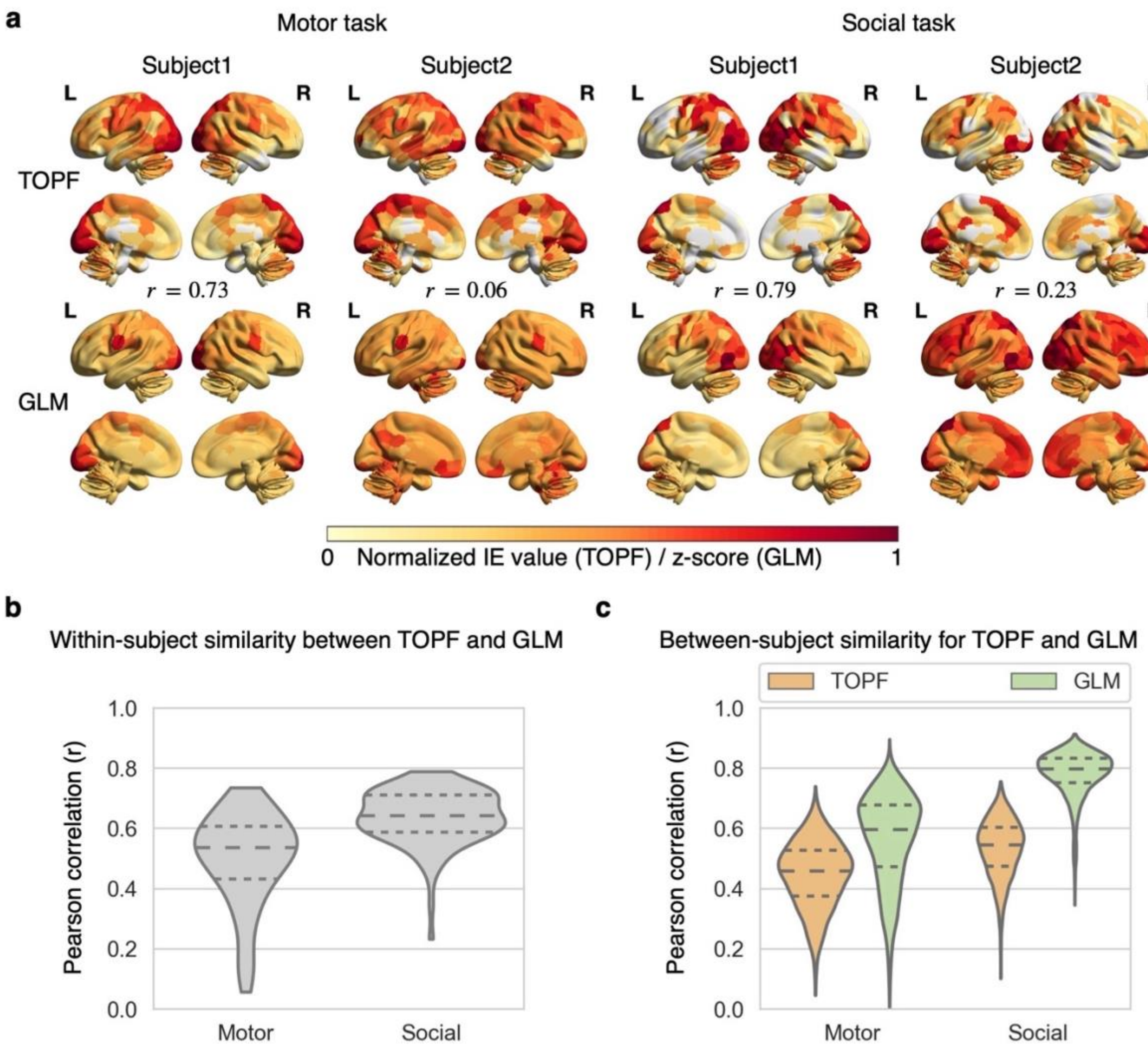


**Supplementary Fig. 1: Validation of TOPF on task-based fMRI data at the individual level. a,** Individual-specific topographies derived by TOPF (upper row) and GLM-derived activation maps (lower row) of two representative subjects for the motor task and social task separately. Each value in the TOPF-identified topographies reflects the IE value (PC1 loading) for the given subject in the given ROI. Each value in the GLM-derived map indicates the activation aggregated across experimental conditions (maximum absolute values of the z-scores) for each subject in the given ROI. For illustration purposes, the IE values and the aggregated z-scores are normalised within each subject to be in [-1,1] and [0,1], respectively, with the IE values being thresholded at 0. The colour from yellow to red indicates the value from low to high. For each task, Subject1 and Subject2 achieve the highest and lowest within-subject similarity (Pearson's correlation coefficient, r) between the TOPF-derived topography and GLM-derived map, respectively. **b,** Distribution of within-subject similarity (r) between the TOPF-derived topography and GLM-derived map over all subjects for the motor and social tasks separately. **c,** Distribution of between-subject similarity (r) over all pairs of subjects for the TOPF-derived topographies and the GLM-derived maps of each task separately. The three dashed lines inside each violin plot from top to bottom denote the third quartile, median and first quartile of the corresponding distribution, respectively.

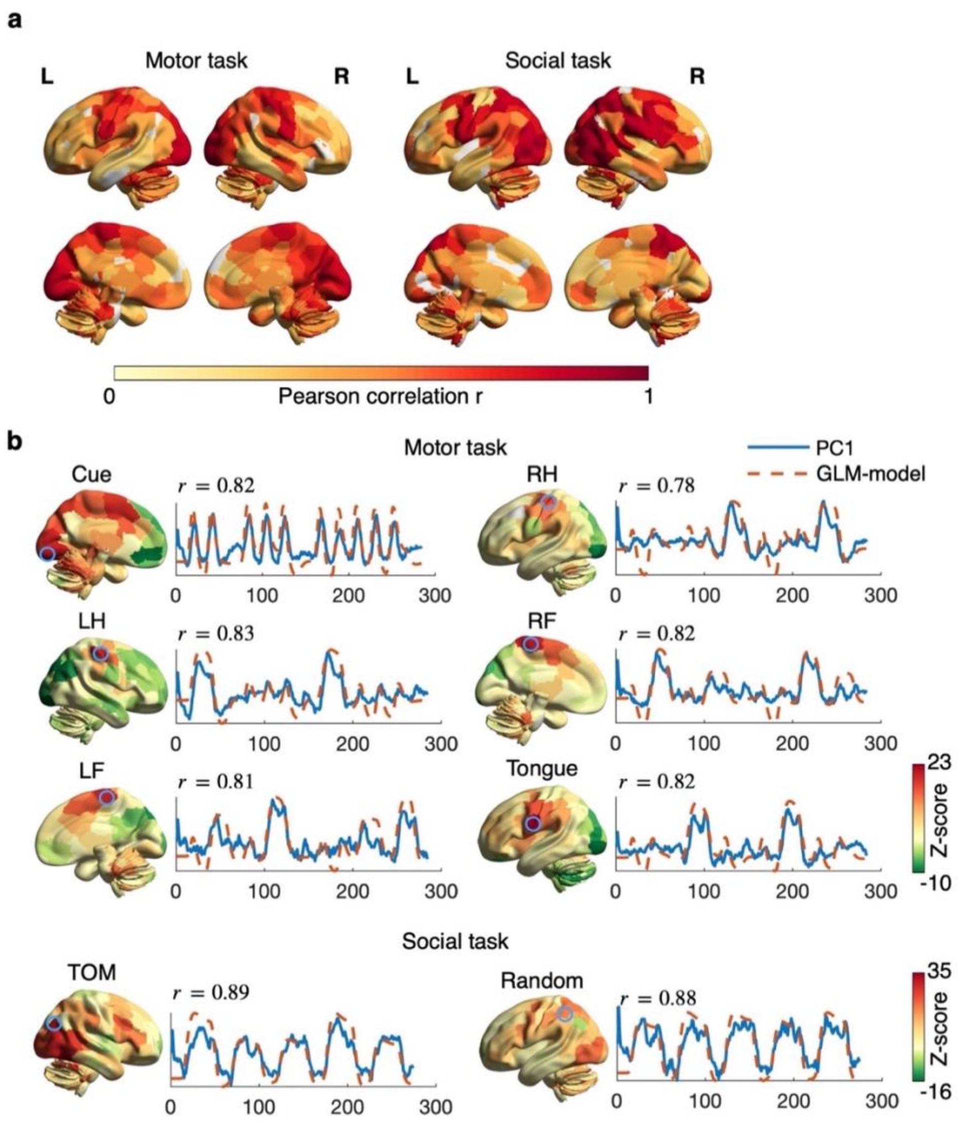


**Supplementary Fig. 2: Validation of TOPF-identified shared response (PC1) time series on task-based fMRI data. a,** Correspondence (Pearson's correlation coefficient, r) between the detected PC1 (blue) and the GLM model (red) of each individual ROI for the motor and social tasks separately. The GLM model is computed as the convolution of HRF with event timing aggregated across all experimental conditions. Only the ROIs for which the correlation is significant ($p<0.05$, Bonferroni corrected) are shown here. The colour from yellow to red indicates the correlation from low to high. **b,** The PC1 time series and the GLM model of a representative ROI for each experimental condition of each task separately. The brain maps are the group-level activation maps derived by GLM for each condition separately, with the colour from green to red indicating the z-score from low to high. These representative ROIs (marked in blue circles) are all strongly activated in the corresponding condition, indicated by the high z-scores in the GLM-derived activation maps. Experimental conditions: Cue: visual cue; LF: left foot; LH: left hand; RF: right foot; RH: right hand; TOM: theory of mind.

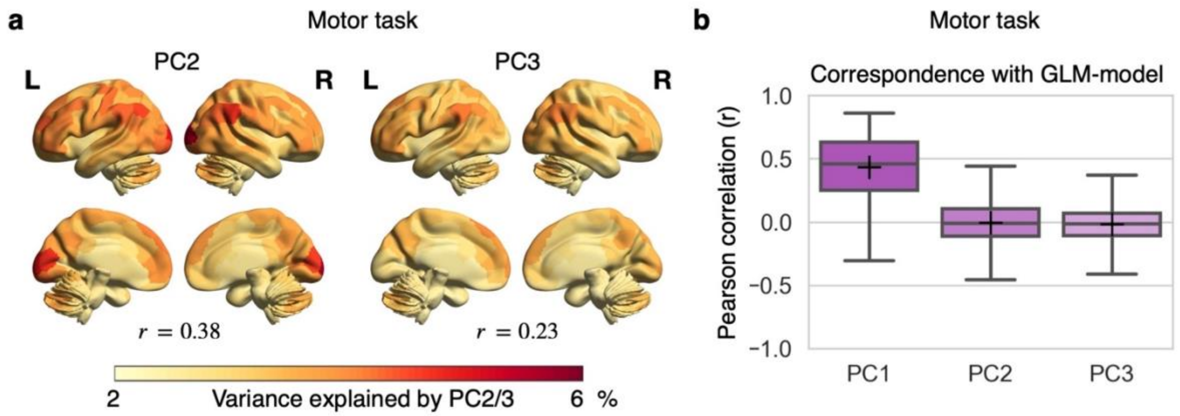


**Supplementary Fig. 3: Comparisons among TOPF-derived PC1, PC2 and PC3 time series in their correspondences with GLM-derived results on task-based fMRI data.** Group-level topographies for PC2 and PC3 separately derived for the motor (**a**) and social tasks (**c**). Each value represents the amount of variance shared across subjects explained by the PC2 or PC3 time series, with the lower and higher values indicated by yellow and red, respectively. The Pearson's correlation (r) between the TOPF-derived topography and the GLM-derived activation map shown in Fig. 2 is computed for each topography of each task separately. Comparisons of PC1, PC2 and PC3 in their correspondence (r) with the model used in GLM (i.e., the convolution of HRF with event timings aggregated across conditions) for the motor (**b**) and social (**d**) tasks. The distribution of the temporal correspondence values over all ROIs is plotted for PC1, PC2 and PC3 separately (box: middle bar, median; upper and lower bars, 75th and 25th percentiles; whisker: upper and lower bars, maximum and minimum; cross: mean).

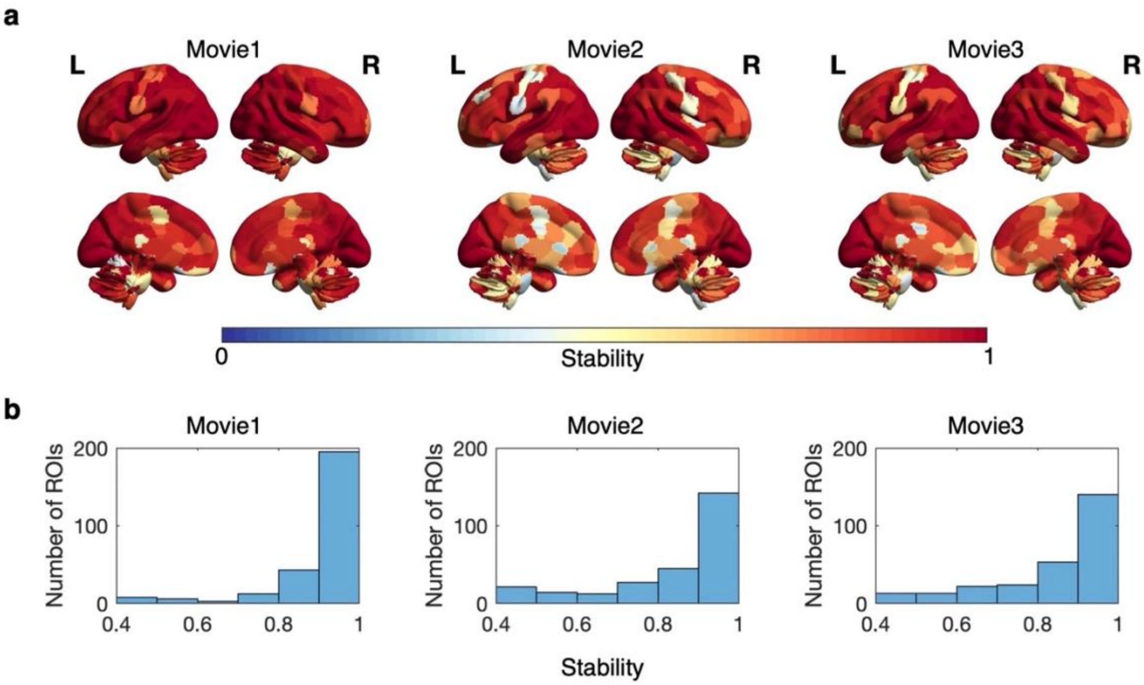


**Supplementary Fig. 4: Stability of TOPF-identified shared response (PC1) time series across 100 subsamples for each movie clip separately on NV fMRI data at n= 90. a,** Stability map of each movie clip. For each ROI, the stability is computed as the mean of the absolute values of the pairwise Pearson's correlation coefficients between subsamples over all pairs of subsamples. The colour from blue to red indicates the stability value from low to high. **b,** Histogram of the stability of all ROIs for each movie clip separately.

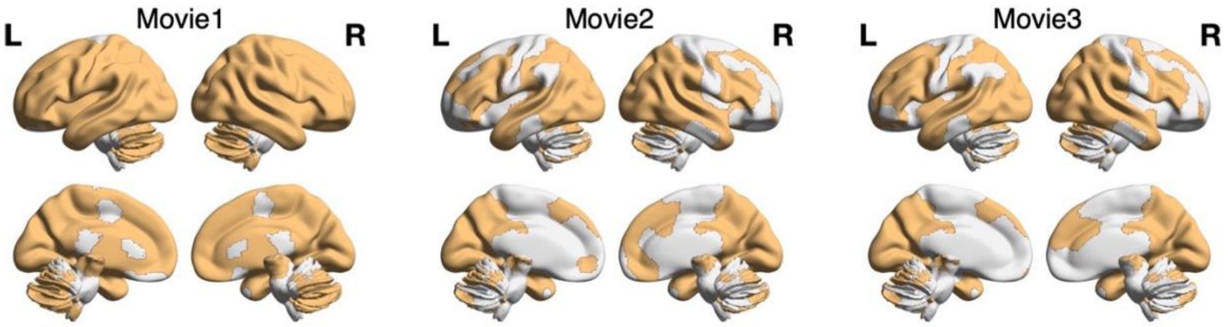


**Supplementary Fig. 5: Statistical significance of the PC1 time series derived by TOPF at n= 90 on NV fMRI data.** Only the ROIs for which the variances explained by the PC1 time series are significantly higher than the chance level (permutation tests with 10000 iterations; Bonferroni-corrected $p<0.05$) consistently on over 80% of all 100 subsamples are shown here.

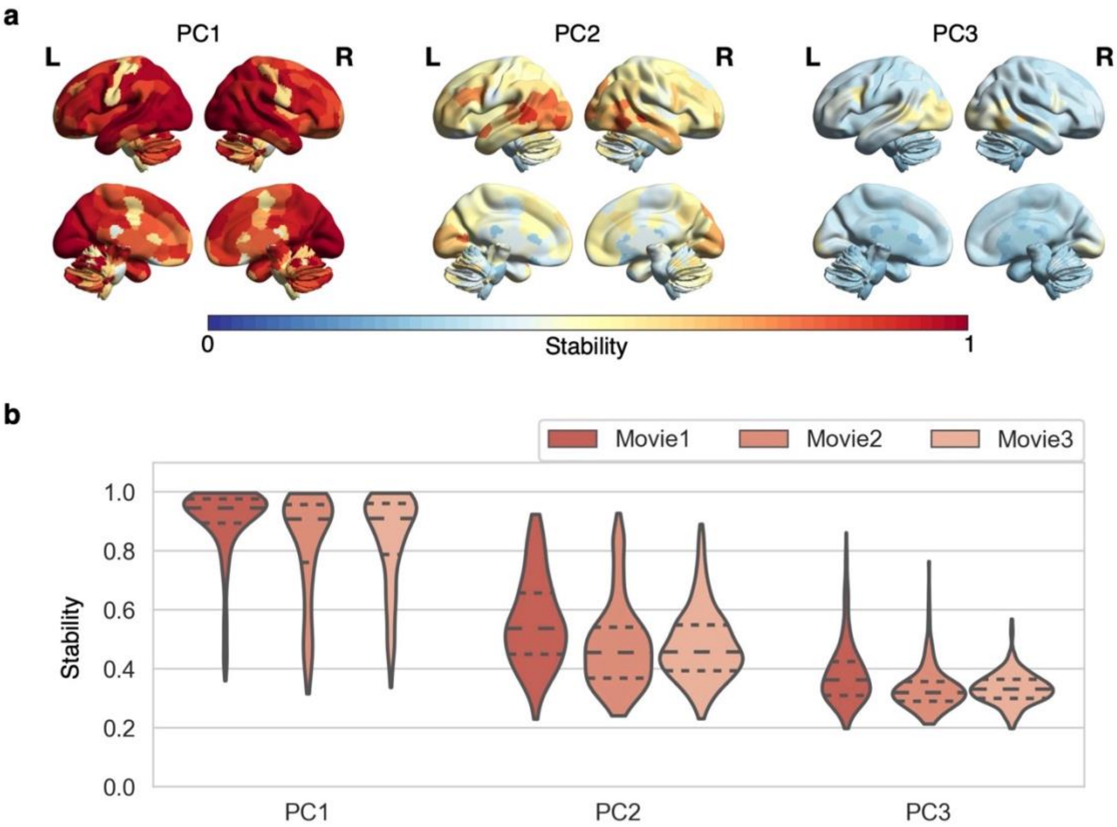


**Supplementary Fig. 6: Comparisons across different PCs in stability at n= 90 on NV fMRI data. a,** Stability map for PC1, PC2 and PC3 separately. Each value in each map denotes the stability of the given ROI averaged over the three movie clips. The colour from blue to red indicates the stability value from low to high. **b,** Distribution of the stability over all ROIs for each PC and each movie clip separately. The three dashed lines inside each violin plot from top to bottom denote the third quartile, median and first quartile of the corresponding distribution, respectively.

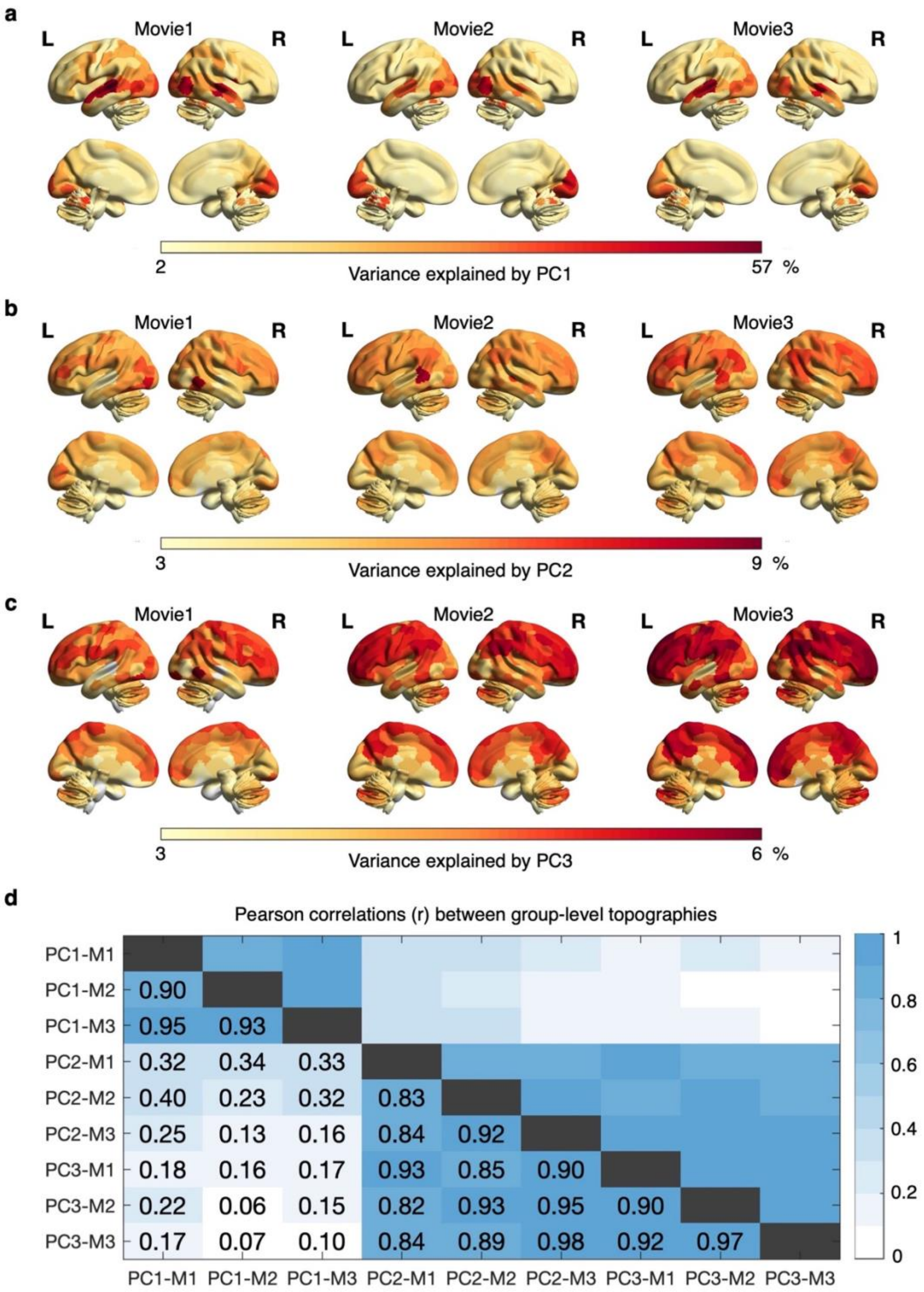


**Supplementary Fig. 7: Comparisons across different PCs in group-level topographies at $n = 90$ on NV fMRI data.** Group-level topography of each movie clip for PC1 (**a**), PC2 (**b**) and PC3 (**c**). Each value in each map denotes the amount of variance explained by the corresponding PC time series in the given ROI aggregated (median) over all 100 subsamples. The colour from yellow to red indicates the value from low to high. Note that the scales are different for different PCs. **d,** Similarity (Pearson's correlation r) between each pair of topographies shown in (a) to (c). M1: Movie1; M2: Movie2; M3: Movie3.

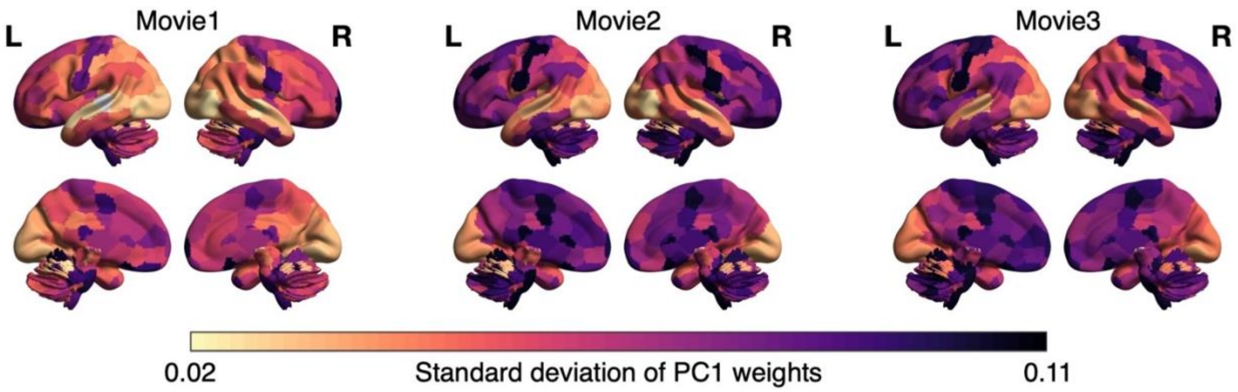


**Supplementary Fig. 8: Brain map of intersubject variability in the IE values derived at n= 90 on NV fMRI data for each movie clip separately**. Each value of each map denotes the intersubject variability in the IE values (standard deviation of PC1 weights across subjects within each subsample averaged over all subsamples) of the given ROI. Note that the PC1 weights (i.e., PC1 loadings normalised by the square root of the eigenvalue of the corresponding PC) are used here instead of the PC1 loadings for fair comparisons across ROIs. The value of intersubject variability from low to high is indicated by the colour from bright to dark.

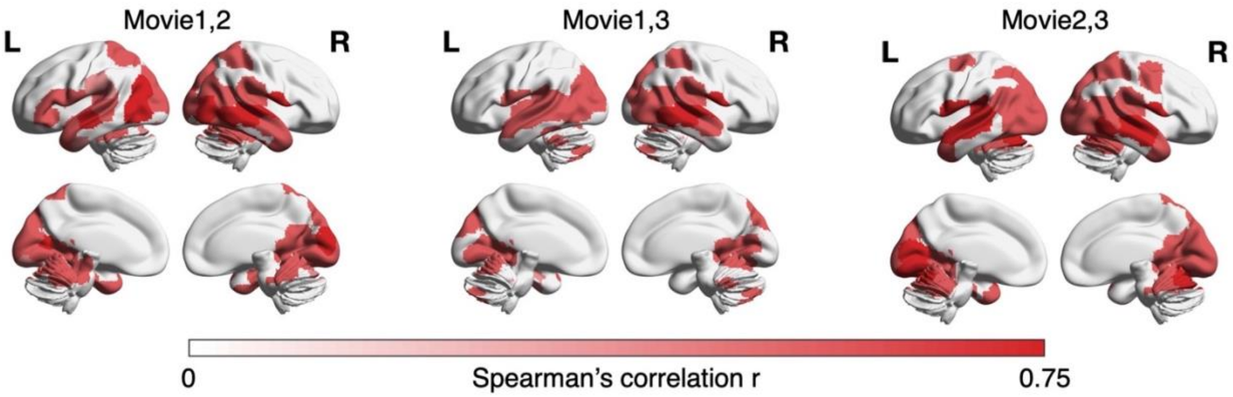


**Supplementary Fig. 9: Similarity in the pattern of the IE values (PC1 loadings) over subjects for each pair of movie clips separately.** For each ROI, the similarity is computed as the Spearman's correlation coefficient (r) of the PC1 loadings (derived at n= 90) over all subjects for each subsample and each pair of movie clips separately. Only the ROIs that consistently achieve a significant correlation (Bonferroni-corrected $p<0.05$) on more than 80% of all 100 subsamples are shown here. For these ROIs, the mean correlation across all subsamples from low to high is indicated by the colour from light to dark.

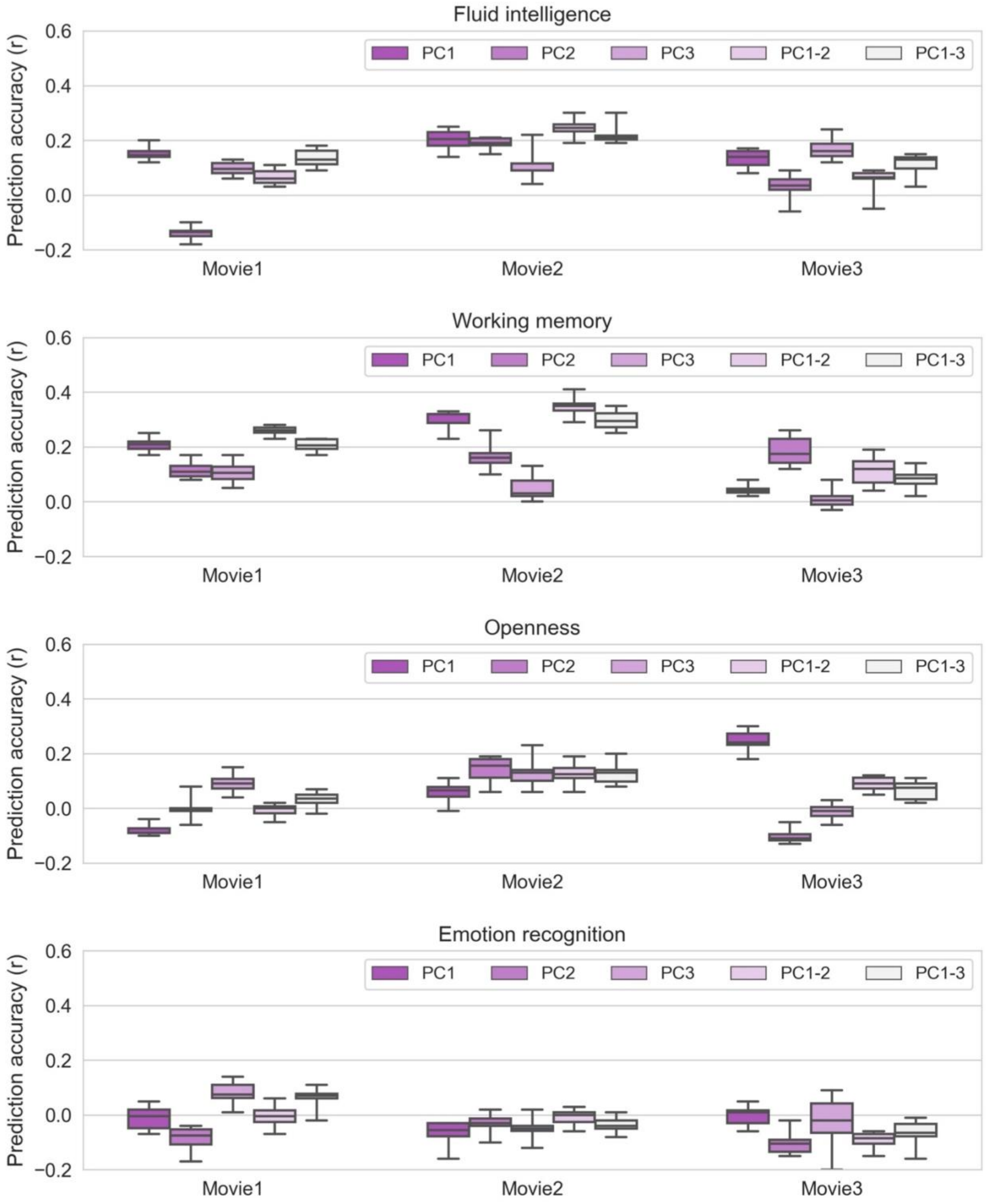


**Supplementary Fig. 10: Comparisons across different PCs in prediction accuracy on NV fMRI data.** For each phenotype, we show the distribution of prediction accuracy (r) over all 10 CV repetitions for each condition and each movie clip separately in a box plot (upper and lower whiskers: maximum and minimum; bars within each box from top to bottom: third quartile, median and first quartile). PC1, PC2 and PC3 represent the conditions where the features used for prediction are the topographies of loadings of PC1, PC2, and PC3, respectively. PC1-2 denotes the condition where we concatenate PC1 and PC2 loadings as features. PC1-3 denotes the condition where we concatenate PC1, PC2 and PC3 loadings as features.

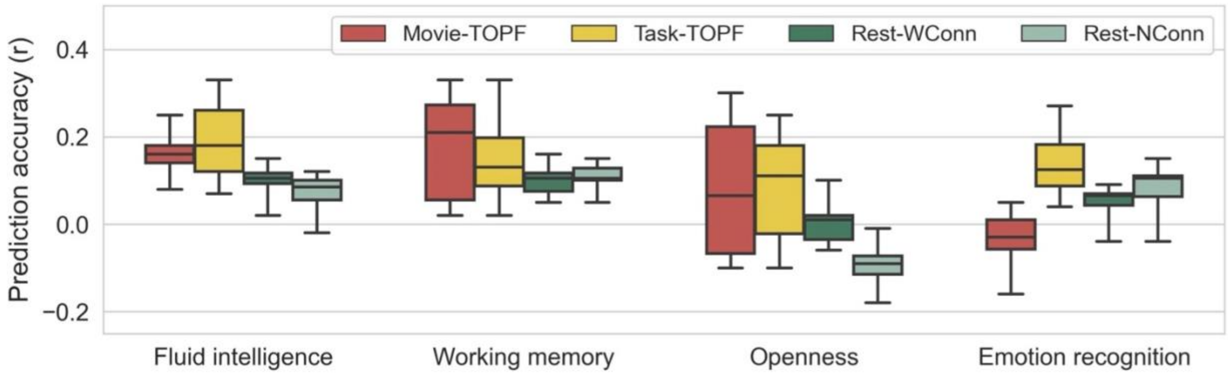


**Supplementary Fig. 11: Comparisons between the performance of TOPF on NV and task fMRI data and the performance of connectivity-based prediction on RS fMRI data.** For each phenotype, we show the distribution of prediction accuracy (r) over all 10 CV repetitions of all the relevant fMRI paradigms for each condition separately. The prediction accuracy (r) is computed in the same way as computed in Fig. 5. Box plot: upper and lower whiskers: maximum and minimum; bars within each box from top to bottom: third quartile, median and first quartile. Movie-TOPF: topographies of PC1 loadings as features on NV fMRI data (the three movie clips); Task-TOPF: topographies of PC1 loadings as features on task fMRI data (the four tasks); Rest-WConn: whole-brain connectomes as features on RS fMRI data; Rest-NConn: nodal connectivity strengths as features on RS fMRI data.

| Stimulus/ Task name | Dataset | Data type | Field strength | # TRs | #Minutes | Stimulus/Task description |
|---|---|---|---|---|---|---|
| Movie1: Two Men | 2 | NV fMRI | 7T | 244 | 04:04 | One man describes his thoughts about him seeing another man keep running; human faces/voices; colour images; background music |
| Movie2: Welcome to Bridgeville | 2 | NV fMRI | 7T | 221 | 03:41 | People talk about themselves living in a small town; human faces/voices; colour images; background music |
| Movie3: Pockets | 2 | NV fMRI | 7T | 189 | 03:08 | People talk about things in their pockets; human faces/voices; colour images; background music |
| Motor | 1, 2 | Task fMRI | 3T | 284 | 03:34 | Movements of body parts; 10 blocks; visual cues; 5 conditions: left/right fingers, left/right toes, tongue |
| Social cognition | 1, 2 | Task fMRI | 3T | 274 | 03:27 | Theory of mind (TOM); 5 blocks; visual stimuli; 2 conditions: mental (TOM) interaction, random interaction |
| Working memory | 2 | Task fMRI | 3T | 405 | 05:01 | N-back tasks; 8 blocks; visual cues; 4 conditions: faces, places, tools, body parts |
| Language | 2 | Task fMRI | 3T | 316 | 03:57 | Story comprehension and verbal math tasks; 8 blocks; auditory stimuli; 2 conditions: story, math |

**Supplementary Table 1: Overview of the included fMRI paradigms of each dataset**. For each paradigm, we list the name of the stimulus or task used during the scan, the dataset(s) it comes from, the fMRI data type, the field strength, the scan duration in volumes (#TR) and in minutes (#Minutes), and the description of the stimulus or task.

| Behaviour/Trait | Domain | Label in HCP | Score range | Score (mean±std) | Behavioural test |
|---|---|---|---|---|---|
| Fluid intelligence | Cognition | PMAT24_A_CR | 6-23 | 16.97±4.42 | Penn Progressive Matrices test |
| Working memory | Cognition | ListSort_Unadj | 80.79-137.98 | 110.09±11.74 | List sorting test; NIH toolbox |
| Openness | Personality | NEOFAC_O | 15-44 | 28.24±5.39 | Personality Five Factor Inventory (NEO-FFI) |
| Agreeableness | Personality | NEOFAC_A | 22-46 | 34.48±5.03 | |
| Conscientiousness | Personality | NEOFAC_C | 17-48 | 34.49±5.74 | |
| Extraversion | Personality | NEOFAC_E | 15-46 | 30.74±6.09 | |
| Neuroticism | Personality | NEOFAC_N | 2-39 | 16.34±6.33 | |
| Emotion recognition | Emotion | ER40_CR | 27-40 | 35.53±2.71 | Emotion recognition test (40 faces) |

**Supplementary Table 2: Overview of the behavioural measures used for phenotype prediction.** For each behavioural measure, we list the name of the behaviour or personal trait it measures, the domain it concerns, the label used by the HCP, the range and the mean with standard deviation (mean±std) of the scores across all subjects, and the test used to obtain the score.

| Confound | Fluid intelligence | Working memory | Openness | Emotion recognition |
|---|---|---|---|---|
| Age | -0.23 | **-0.28** | -0.11 | 0.05 |
| Sex | 0.15 | **0.26** | 0.21 | -0.05 |
| HM-Movie1 | -0.23 | -0.16 | -0.03 | **-0.30** |
| HM-Movie2 | -0.21 | -0.15 | -0.02 | **-0.27** |
| HM-Movie3 | -0.18 | -0.13 | -0.08 | **-0.30** |
| HM-Motor | **-0.26** | **-0.26** | -0.06 | **-0.39** |
| HM-Social | **-0.25** | -0.16 | -0.06 | **-0.31** |
| HM-WM | **-0.28** | **-0.24** | -0.02 | **-0.27** |
| HM-Language | -0.20 | **-0.24** | -0.06 | **-0.24** |
| HM-RS | -0.20 | -0.07 | -0.07 | **-0.29** |

**Supplementary Table 3: Correlations between phenotypes and confounding variables.** The Pearson's correlation coefficient between each phenotype and each of the three confounding variables, including age, sex and head motion (HM), is computed. HM is measured for each fMRI paradigm separately as the relative root-mean-square framewise displacement. Significant results (Bonferroni-corrected $p<0.05$) are highlighted in bold. WM: working memory task; RS: resting state.

| | Movie1 | | Movie2 | | Movie3 | |
|---|---|---|---|---|---|---|
| **Phenotypes** | **r** | **p-value** | **r** | **p-value** | **r** | **p-value** |
| Fluid intelligence | **0.15** | 0.025 | **0.20** | 0.009 | 0.13 | 0.056 |
| Working memory | **0.21** | 0.005 | **0.30** | 0 | 0.04 | 0.293 |
| Emotion recognition | -0.01 | 0.532 | -0.06 | 0.769 | -0.00 | 0.496 |
| Openness | -0.08 | 0.849 | 0.06 | 0.245 | **0.24** | 0.002 |
| Agreeableness | -0.03 | 0.63 | -0.06 | 0.753 | -0.07 | 0.802 |
| Conscientiousness | 0.14 | 0.05 | -0.13 | 0.929 | 0.02 | 0.406 |
| Extraversion | 0.03 | 0.358 | -0.02 | 0.585 | -0.03 | 0.618 |
| Neuroticism | -0.08 | 0.837 | -0.11 | 0.904 | -0.12 | 0.926 |

**Supplementary Table 4: Prediction performance of TOPF on NV fMRI data for all 8 phenotypes in a 10-fold CV with 10 repetitions.** The accuracy is computed in the same way as computed in Fig. 5, where the Pearson's correlation coefficient (r) between predicted and observed scores is computed after regressing out the three confounding variables (age, sex and head motion) from both scores. For each fMRI paradigm-phenotype combination, we show the mean accuracy as well as the mean p-value (permutation tests with 5000 iterations) over the 10 repetitions. Significant results ($p<0.05$) are highlighted in bold.

| | Motor | | Social | | WM | | Language | |
|---|---|---|---|---|---|---|---|---|
| **Phenotypes** | **r** | **p-value** | **r** | **p-value** | **r** | **p-value** | **r** | **p-value** |
| Fluid intelligence | 0.13 | 0.056 | **0.23** | 0.003 | 0.11 | 0.088 | **0.27** | 0.001 |
| Working memory | 0.07 | 0.19 | 0.13 | 0.056 | 0.11 | 0.085 | **0.27** | 0.001 |
| Emotion recognition | 0.11 | 0.089 | **0.23** | 0.002 | 0.1 | 0.124 | 0.11 | 0.089 |
| Openness | 0.04 | 0.301 | **0.14** | 0.038 | -0.07 | 0.815 | **0.22** | 0.004 |
| Agreeableness | 0.12 | 0.078 | 0.1 | 0.125 | 0.01 | 0.474 | 0.04 | 0.328 |
| Conscientiousness | 0.13 | 0.059 | 0.01 | 0.47 | -0.07 | 0.811 | -0.04 | 0.693 |
| Extraversion | -0.02 | 0.603 | -0.07 | 0.773 | 0.04 | 0.332 | 0.03 | 0.367 |
| Neuroticism | -0.13 | 0.947 | -0.13 | 0.948 | 0.02 | 0.419 | 0.12 | 0.065 |

**Supplementary Table 5: Prediction performance of TOPF on task-based fMRI data for all 8 phenotypes in a 10-fold CV with 10 repetitions.** The accuracy is computed in the same way as computed in Fig. 5, where the Pearson's correlation coefficient (r) between predicted and observed scores is computed after regressing out the three confounding variables (age, sex and head motion) from both scores. For each fMRI paradigm-phenotype combination, we show the mean accuracy as well as the mean p-value (permutation tests with 5000 iterations) over the 10 repetitions. Significant results ($p<0.05$) are highlighted in bold.

| Phenotypes | Movie1 | Movie2 | Movie3 | Motor | Social | WM | Language |
|---|---|---|---|---|---|---|---|
| Fluid intelligence | 0.21 | 0.26 | 0.12 | 0.20 | 0.29 | 0.19 | 0.31 |
| Working memory | 0.22 | 0.33 | 0.05 | 0.13 | 0.22 | 0.19 | 0.34 |
| Emotion recognition | 0.02 | -0.04 | 0.01 | 0.17 | 0.28 | 0.11 | 0.12 |
| Openness | -0.07 | 0.07 | 0.23 | 0.04 | 0.16 | -0.07 | 0.22 |
| Agreeableness | -0.01 | -0.06 | -0.05 | 0.13 | 0.12 | 0.01 | 0.08 |
| Conscientiousness | 0.15 | -0.13 | 0.02 | 0.13 | 0.00 | -0.07 | -0.05 |
| Extraversion | 0.04 | -0.02 | -0.03 | -0.01 | -0.06 | 0.03 | 0.02 |
| Neuroticism | -0.08 | -0.1 | -0.12 | -0.11 | -0.12 | 0.04 | 0.14 |

**Supplementary Table 6: Prediction accuracy of TOPF in a 10-fold CV with 10 repetitions without confound removal.** For each fMRI paradigm-phenotype combination, we show the mean accuracy (the Pearson's correlation coefficient r between predicted and observed scores across all subjects without confound removal) over the 10 repetitions.

| **Phenotypes** | **Movie1** | **Movie2** | **Movie3** | **Motor** | **Social** | **WM** | **Language** |
|---|---|---|---|---|---|---|---|
| Fluid intelligence | 0.16 | 0.19 | 0.15 | 0.15 | **0.28** | 0.1 | **0.30** |
| Working memory | **0.22** | **0.30** | 0.03 | 0.07 | 0.14 | 0.14 | **0.30** |
| Emotion recognition | -0.02 | -0.08 | 0.01 | 0.16 | **0.25** | 0.09 | 0.09 |
| Openness | -0.10 | 0.06 | **0.29** | 0.04 | 0.13 | -0.07 | **0.23** |
| Agreeableness | -0.03 | -0.04 | -0.06 | 0.12 | 0.13 | -0.01 | 0.05 |
| Conscientiousness | 0.14 | -0.14 | 0.02 | 0.12 | 0.02 | -0.08 | -0.06 |
| Extraversion | 0.02 | -0.01 | -0.02 | 0.01 | -0.07 | 0.05 | 0.04 |
| Neuroticism | -0.05 | -0.11 | -0.10 | -0.16 | -0.12 | 0.03 | 0.12 |

**Supplementary Table 7: Prediction accuracy of TOPF in a leave-one-out CV.** For each fMRI paradigm-phenotype combination, the accuracy is computed as the Pearson's correlation coefficient (r) between predicted and observed scores over all subjects after regressing out the three confounding variables (age, sex and head motion) from both scores. Significant results (FDR corrected $p<0.05$; permutation tests with 5000 iterations) are highlighted in bold.
